# Effect of continuous compressive force on the expression of RANKL, OPG, and VEGF in MC3T3-E1 and MLO-Y4 cells

**DOI:** 10.1101/474254

**Authors:** Yuka Yashima, Masato Kaku, Taeko Yamamoto, Jin Izumino, Haruka Kagawa, Kotaro Tanimoto

## Abstract

Osteocytes, known to have mechano-sensory functions, influence the regulation of bone remodeling. However, the mechanism by which osteocytes regulate bone metabolism when mechanical forces are being applied is still unclear. Osteoclastogenesis is mainly regulated by receptor activator of nuclear factor kappa-B ligand (RANKL); the protein osteoprotegerin (OPG) and angiogenesis also play important roles in osteogenesis. RANKL, OPG, and vascular endothelial growth factor (VEGF) are thought to be key factors for bone metabolism. In this study, we examined the effect of a continuous compressive force (CF) on the expression of RANKL, OPG, and VEGF in osteoblastic murine osteocytes (MLO-Y4) and osteoblastic (MC3T3-E1) cells. Gene and protein expression levels of RANKL, OPG, and VEGF in MLO-Y4 and MC3T3-E1 cells were quantitatively determined by real-time PCR and enzyme-linked immunosorbent assay (ELISA). Both cell types were also subjected to a CF of 1.0 g/cm^2^ for 1, 3, 6, and 12 hours. Furthermore, the effect of a stretch-activated (S-A) channel was examined by gadolinium (Gd^3+^) administration. The ratio of gene and protein expressions of RANKL, VEGF, and RANKL/OPG in MLO-Y4 cells were significantly higher than in MC3T3-E1 cells, while the expression of OPG was significantly lower. After CF application, both cell types showed significant increases in RANKL and VEGF expression as well as the RANKL/OPG ratio. Additionally, the upregulated gene and protein levels of these factors were reduced by Gd^3+^ administration.

These findings suggest that osteocytes play more important roles in bone metabolism and angiogenesis than osteoblasts. Osteocytes regulate the expression of RANKL, OPG, and VEGF via the S-A channel through the response to mechanical stress.

## Introduction

Bone tissue shows constant remodeling by osteoblastic bone formation and osteoclastic bone resorption in order to maintain bone volume and homeostasis [1, 2]. Therefore, bone remodeling has been suggested to be regulated by a crosstalk between osteoblasts, osteoclasts, and osteocytes [3]. Receptor activator of nuclear factor kappa-B ligand (RANKL) [4] and osteoprotegerin (OPG) [5] are crucial factors for osteoclast differentiation. Osteoblasts have been thought to be the main source of RANKL for osteoclastogenesis [6, 7], and therefore investigations of RANKL expression and their mechanism have been mostly studied in osteoblastic cells [8, 9]. Vascular endothelial growth factor (VEGF), which is a mitogen specific for vascular endothelial cells, plays a major role in angiogenesis [10]. As bone tissue is rich in blood vessels, and bone remodeling requires neovascularization, VEGF is thought to be an important factor not only for angiogenesis but also for skeletal development and bone regeneration [11].

It is well known that mechanical stress can influence the regulation of bone remodeling. Nettelhoff et al. reported that the highest RANKL/OPG ratio was observed after the application of 5% compressive force (CF) in osteoblasts [12]. Tripuwabhrut et al. also suggested that CFs on osteoblasts enhance osteoclastogenesis by an increase in RANKL expression and decrease in OPG expression [13]. It has been reported that cyclic tensile forces increase the mRNA and protein expressions of VEGF in osteoblastic MC3T3-E1 cells [14], and this reaction was inhibited by gadolinium (Gd^3+^), a stretch-activated (S-A) channel blocker [15]. Reher et al. showed that the production of VEGF in human osteoblasts can be increased by ultrasounds at 1 MHz and 45 kHz [16]. These results suggest that the application of mechanical stress on osteoblasts can affect the expressions of RANKL, OPG, and VEGF, thereby accelerating bone remodeling.

Osteocytes, which account for more than 90% of the cells in bone tissue, are derived from osteoblasts and are embedded in the bone matrix and function as mechanosensory cells through the lacuno-canalicular network [17]. Recently, it was shown that osteocytes can be a main source of RANKL during bone remodeling [18, 19]. However, the effects of mechanical forces on the expression of factors related to bone remodeling between osteocytes and osteoblasts are still unclear. In this study, we examined the effects of CF on the expression of RANKL, OPG, and VEGF in murine osteocytes (MLO-Y4) and compared to that in osteoblastic (MC3T3-E1) cells. We also investigated the influence of Gd^3+^, which blocks the S-A channel upon the expression of RANKL, OPG, and VEGF.

## Materials and methods

### Cell culture

Murine osteoblastic MC3T3-E1 cells were obtained from RIKEN Cell Bank (Tsukuba, Japan), and cultured in alpha minimum essential medium (α-MEM) supplemented with 10% FBS. All cell lines were cultured with 240 ng/mL kanamycin (Meiji Seika, Tokyo, Japan), 1 mg/mL amphotericin-B (ICN Biomedicals Corp, Costa Mesa, CA, USA), 500ng/mL penicillin (Sigma Aldrich, Saint Louis, MO, USA) at 37°C in 5% CO_2_. Murine osteocyte-like cells (MLO-Y4 cells) were obtained from Kerafast (Boston, MA, USA), and cultured on collagen-coated plates (pig tendon type I collagen, IWAKI, Tokyo, Japan) in α-MEM (Sigma Aldrich, Saint Louis, MO, USA) with 2.5% heat inactivated fetal bovine serum (FBS, Daiichi Chemical, Tokyo, Japan) and 2.5% heat inactivated newborn calf serum (CS, Thermo Fisher Scientific, Waltham, MO, USA). The medium was changed twice a week, and cells were subcultured by treatment with 0.05% trypsin and 0.53 mM ethylenediaminetetraacetic acid (EDTA) followed by platting at a density of 5×10^4^ cells/well in 6-well plates. For all experiments, cells between the 4th and the 6th passages were used.

### Application of CF

MC3T3-E1 and MLO-Y4 cells were continuously compressed by a uniform compression method with serum-free conditioned media in 6-well plates at a density of 50,000 cells according to the previous study by Tripuwabhrut et al. (Fig. 1) [20]. A thin glass plate was placed over a confluent cell layer, and the cells were subjected a CF of 1.0 g / cm^2^ for various loading time (1, 3, 6, and 12 hours) to examine the expressions of RANKL, OPG, VEGF, and RANKL/OPG ratio.

**Fig 1.**
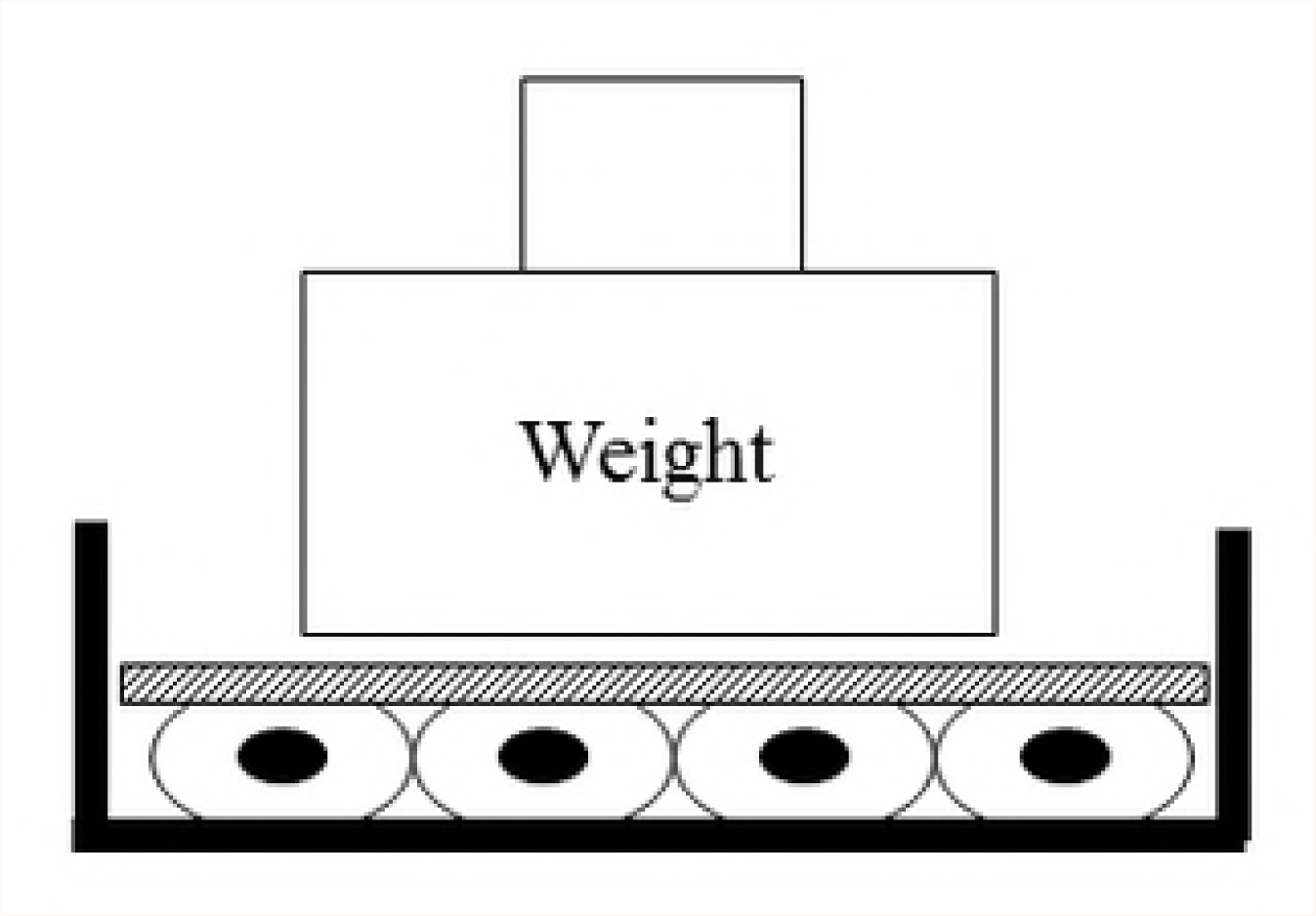
A schematic drawing of compression force application (CF) on the cell layer. 1.0 g / cm^2^ weight is placed over a confluent cell layer.

### Total RNA extraction and cDNA synthesis

Total RNA was isolated from the cell cultures with or without the application of CF using a Quickprep Total RNA extraction kit (Amersham Biosciences, Tokyo, Japan). Single-stranded cDNA was synthesized from 1 μg of total RNA using Oligo (dT)_20_ primer (Toyobo, Osaka, Japan) and a Rever Tra Ace-α first-strand cDNA synthesis kit (Toyobo).

### Primers

RANKL: 5’- CATCGCTCTGTTCCTGTACTTTC -3’ (forward),

5’- AGGAGTCAGGTAGTGTGTCTTCA -3’ (reverse);

OPG: 5’- ACCCAGAAACTGGTCATCAGC -3’ (forward),

5’- CTGCAATACACACACTCATCACT -3’ (reverse);

VEGF: 5’- ATGCGGATCAAACCTCA -3’ (forward),

5’- TTCTGGCTTTGTTCTGTCTT -3’ (reverse);

glyceraldehyde-3-phosphate dehydrogenase (G3PDH) primers (Rever Tra Ace-α,

First-strand cDNA Synthesis Kit, Toyobo) was used as a control primer:

5’- ATGGCCTTCCGTGTTCCT -3’ (forward),

5’- CCCAAGATGCCCTTCAGT -3’ (reverse).

### Quantitative real-time polymerase chain reaction (PCR) analysis

Quantitative real-time PCR was carried out using the SYBR Green I assay in conjunction with an ABI Prism 7700 sequence detection system (Biosystems, Foster City, CA, USA). A template cDNA at a volume of 1 μL was used during the PCR under the following parameters: 2 min at 50°C; 10 min at 95°C; and then 40 cycles of 45 sec at 94°C, 45 sec at 55°C, and 45 sec at 72°C. SYBER Green I dye intercalation into the minor groove of double-stranded DNA reached maximum emission at 530 nm. PCR reactions for each sample were repeated three times for both the target gene and the control. Quantitative results of real-time fluorescence PCR were assessed by a cycle threshold (Ct) value, which identifies a cycle when the fluorescence of a given sample becomes significantly different from the baseline signal. Relative quantifications of the RANKL, OPG, and VEGF signals were normalized and expressed relative to the G3PDH signals.

### Measurement of protein concentrations of RANKL, OPG, VEGF, and RANKL/OPG ratio

The culture medium with or without the application of CF was collected and cleared at 3000 rpm for 5 minutes for enzyme-linked immunosorbent assay (ELISA). The amount of protein concentration of RANKL (Murine sRANK Ligand Mini ABTS ELISA Development kit, PeproTech Inc, Rocky Hill, NJ, USA), OPG (Osteoprotegerin Mouse Immunoassay kit, R&D Systems, Minneapolis, MN, USA), and VEGF (Murine VEGF Mini ABTS ELISA Development kit, PeproTech Inc, Rocky Hill, NJ, USA) were measured using the quantitative sandwich enzyme immunoassay technique according to the manufacturer’s instructions. Standard curves were obtained as usual, and the experiment was repeated three times.

### Effects of Gadolinium (Gd^3+^) on the expressions of RANKL, OPG, and VEGF as well as the RANKL/OPG ratio in MLO-Y4 cells

The effects of Gd^3+^ on the expression of RANKL, OPG, and VEGF in MLO-Y4 cells were examined under CF group. Because it has been reported that 1–100 μM gadolinium inhibited S-A channels [21], cells were incubated with 10 μM Gd^3+^ chloride hexahydrate (Wako, Osaka, Japan) for 30 min. After treatment of Gd^3+^, 1.0 g/cm^2^ CF was applied for 3 hours to assess the effects of CF on the expressions of RANKL, OPG, VEGF mRNAs and RANKL/OPG ratio. Furthermore, 1.0 g/cm^2^ CF was also applied for 12 hours to examine the amounts of RANKL, OPG, VEGF proteins and RANKL/OPG ratio.

### Statistical treatment

The Student’s *t*-test was used to evaluate statistical differences in RANKL, OPG, and VEGF mRNA and protein expressions as well as the RANKL/OPG ratio between the MLO-Y4 and MC3T3-E1 cells. Statistical significances in mRNA and protein levels after the application of CF was assessed by analysis of variance followed by the Fisher’s method. A p < 0.05 was considered statistically significant.

## Results

### Expression of mRNA and protein concentrations of RANKL, OPG, VEGF, and RANKL/OPG ratio in MT3T3-E1 and MLO-Y4 cells

RANKL and VEGF mRNA expressions and the RANKL/OPG ratio in MLO-Y4 cells were significantly higher than that in MC3T3-E1 cells (Figs. 2A, 2C, and 2D). The gene expression level of OPG was significantly lower in MLO-Y4 cells than that in MC3T3-E1 cells (Fig. 2B).

**Figs 2.**
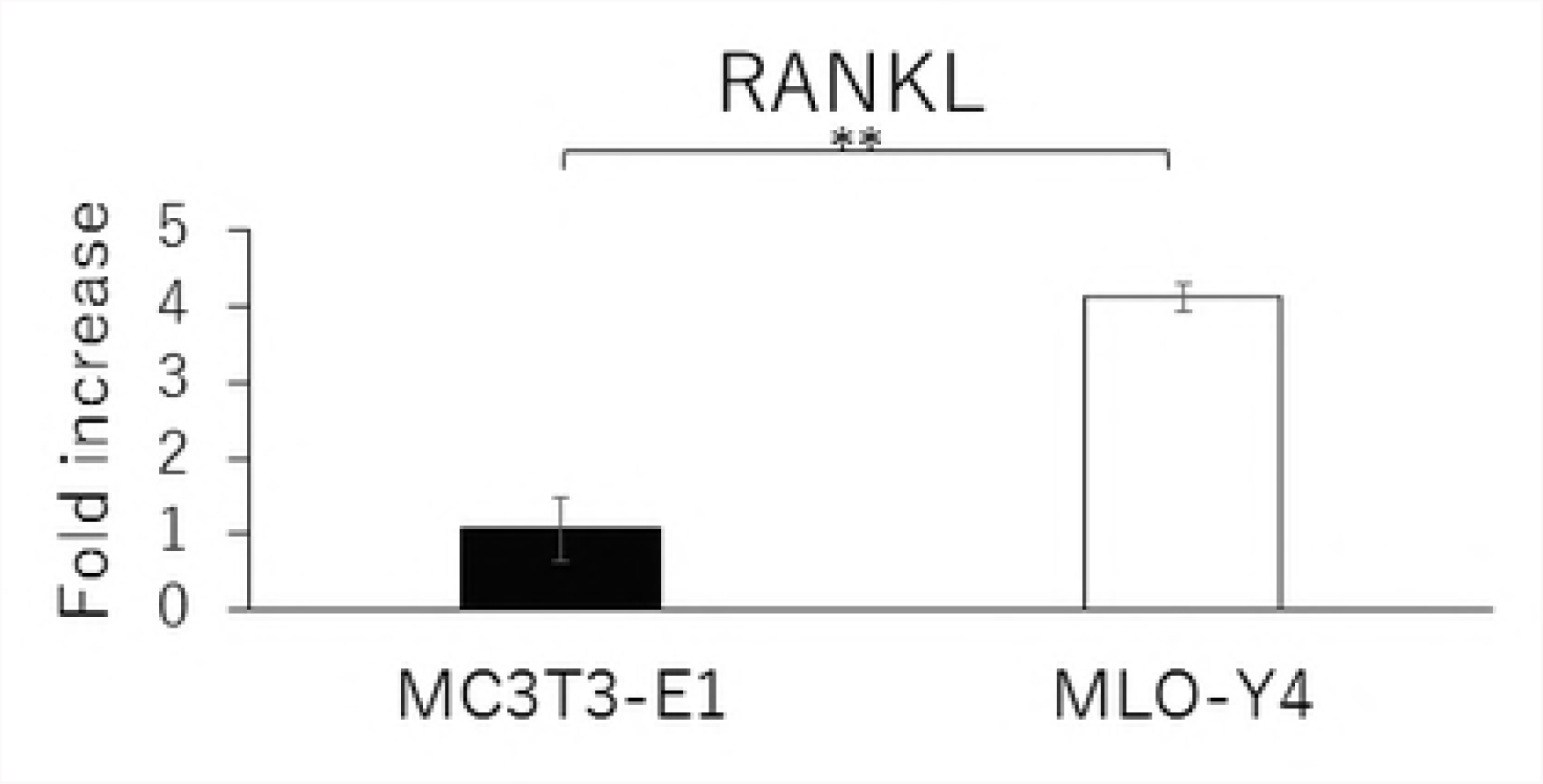
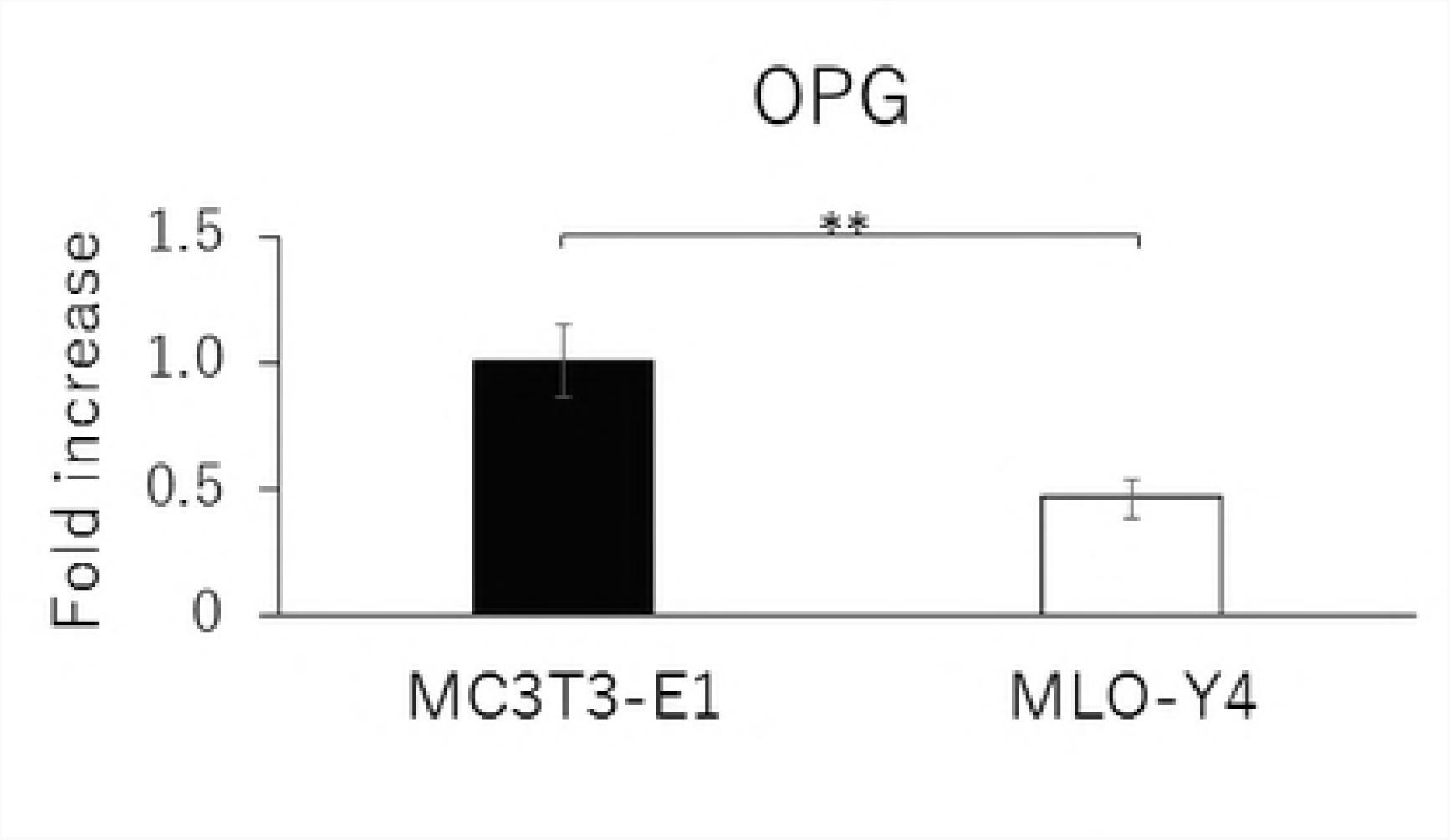
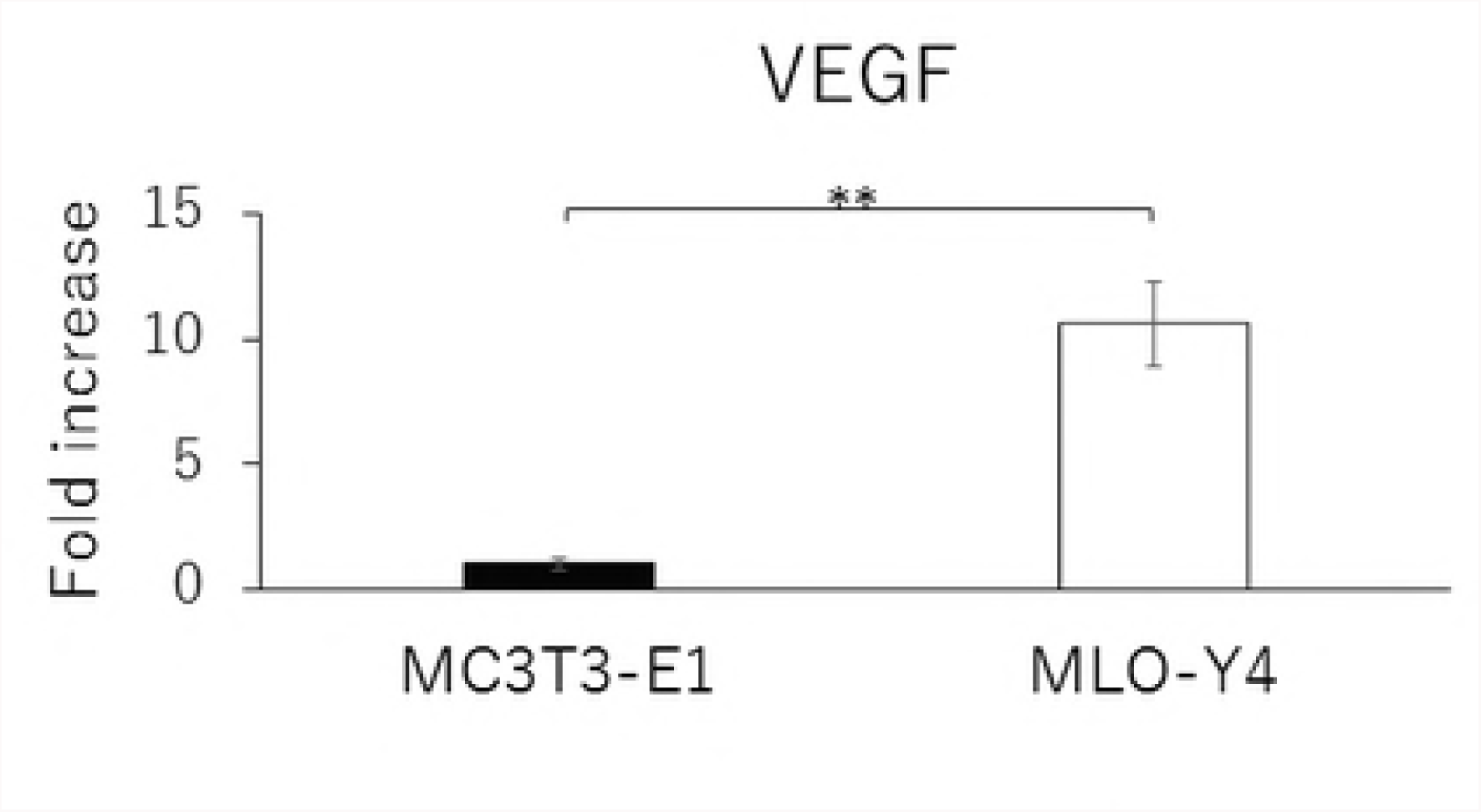
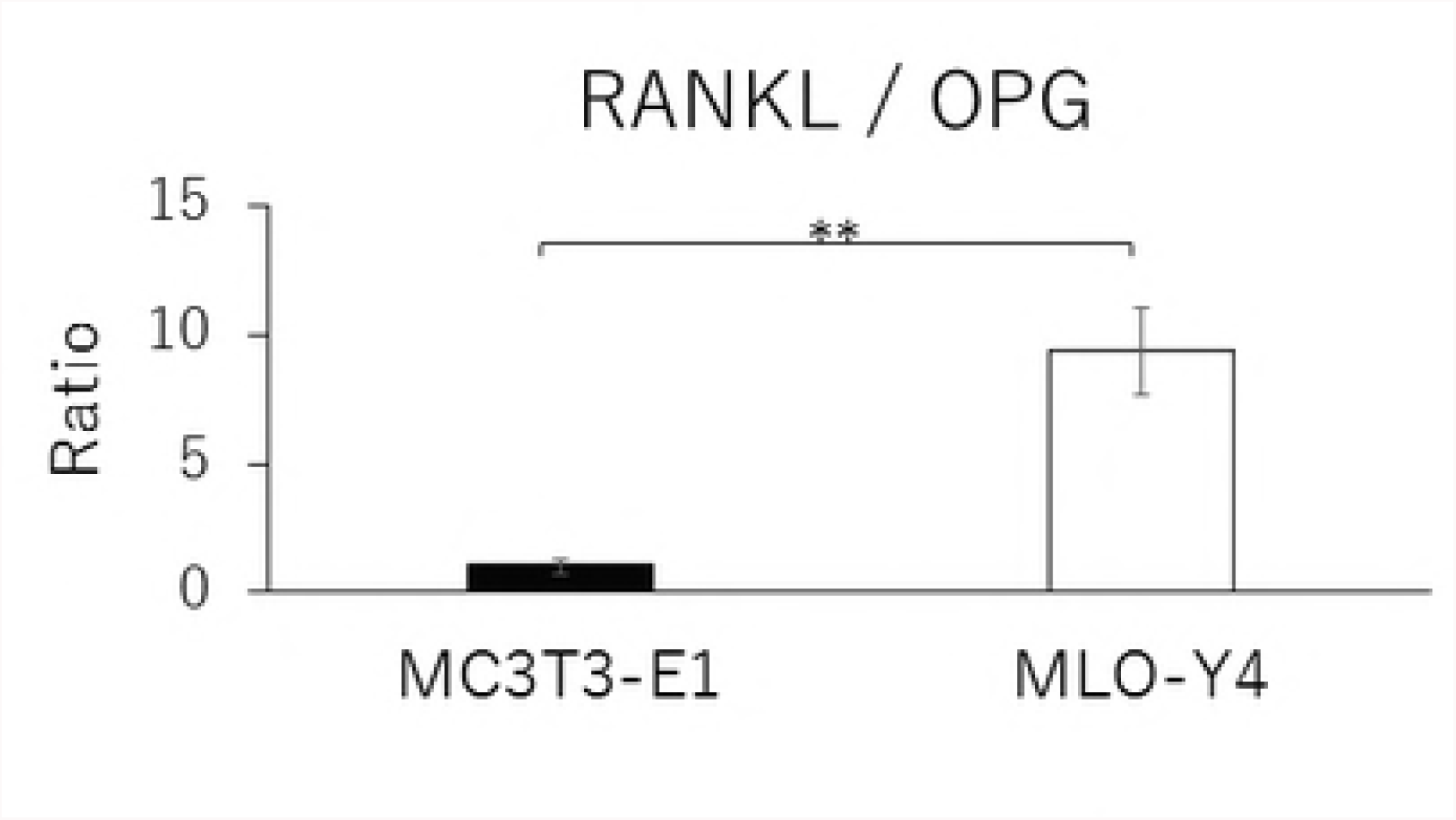
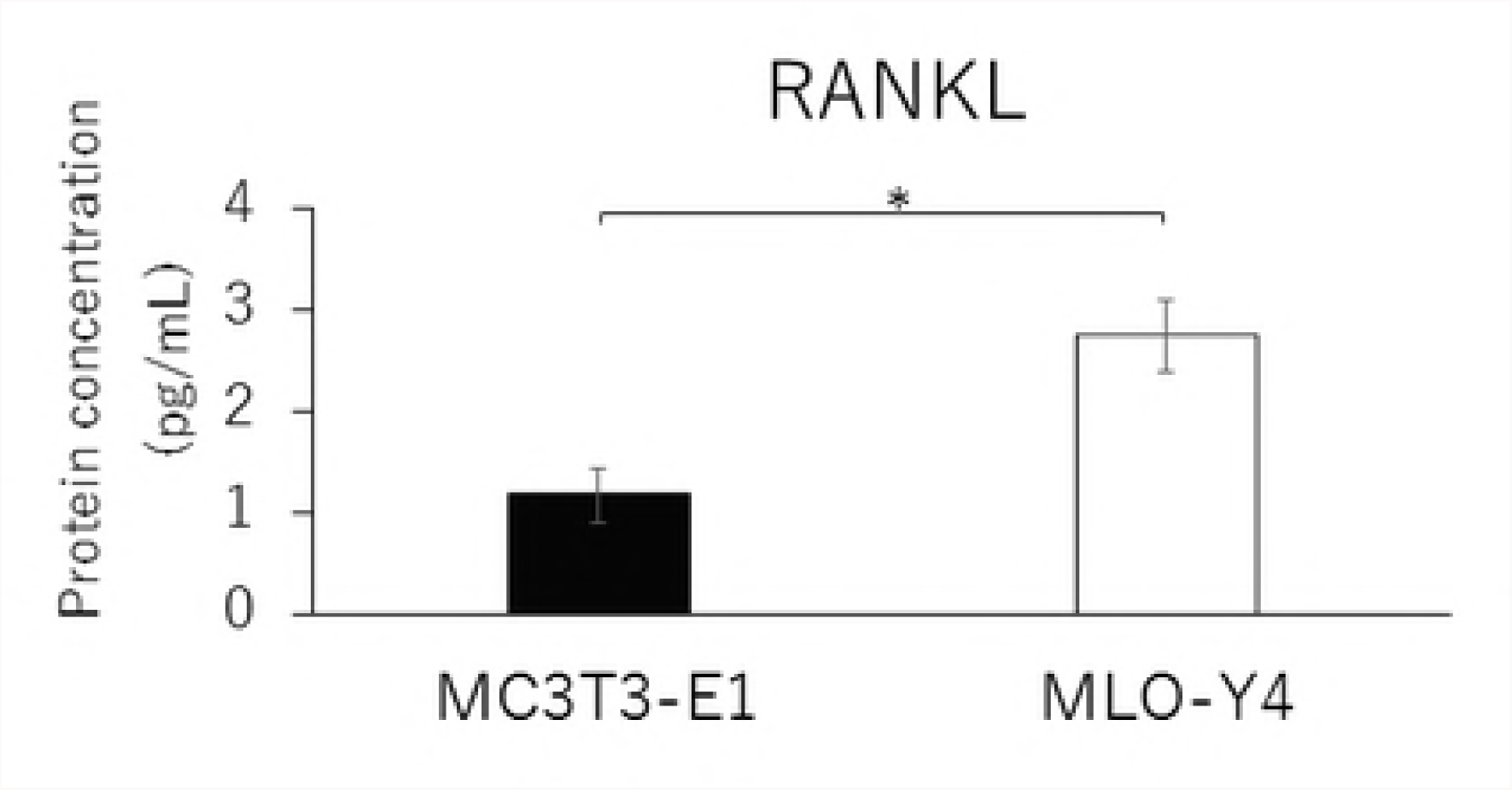
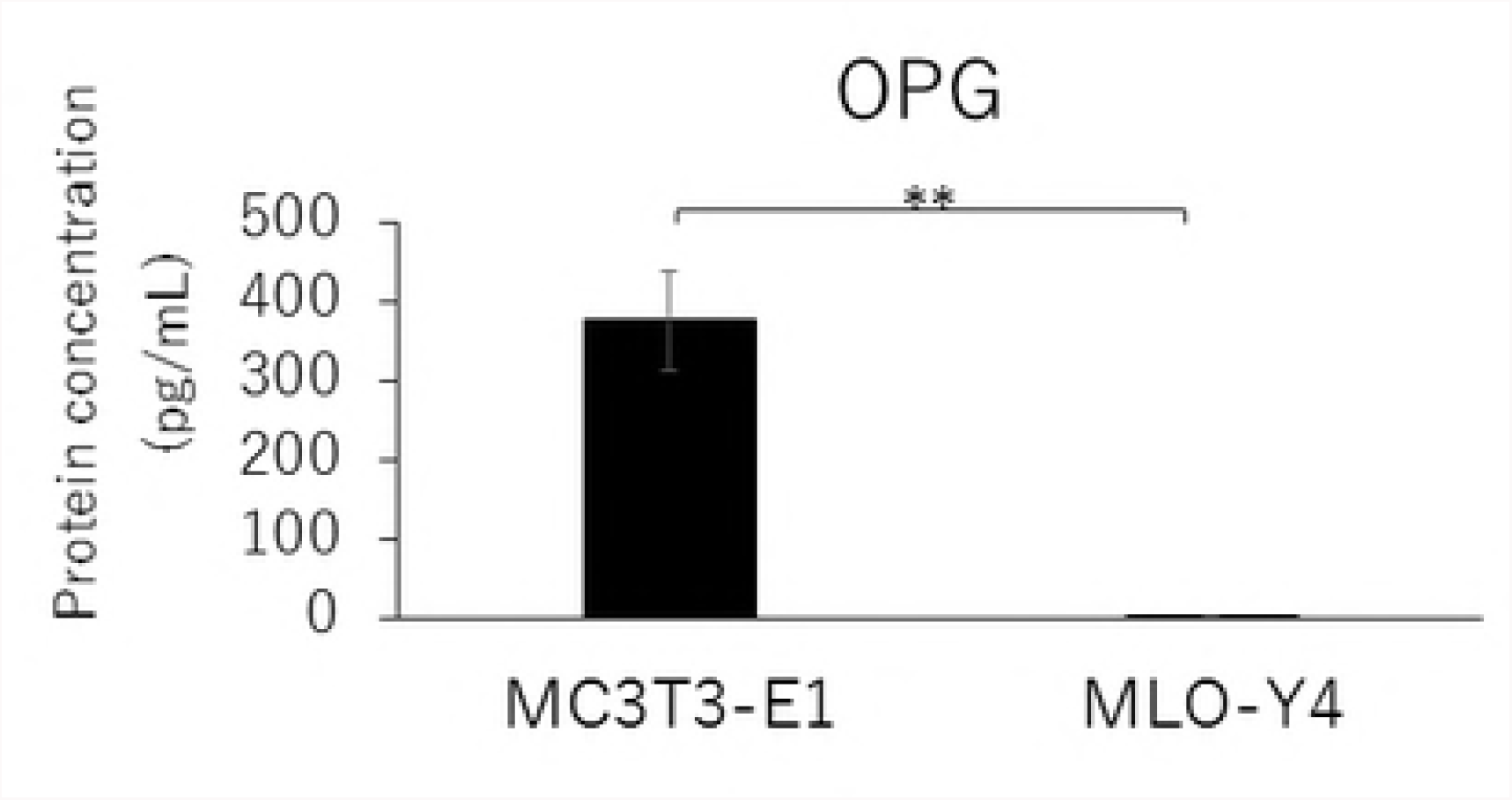
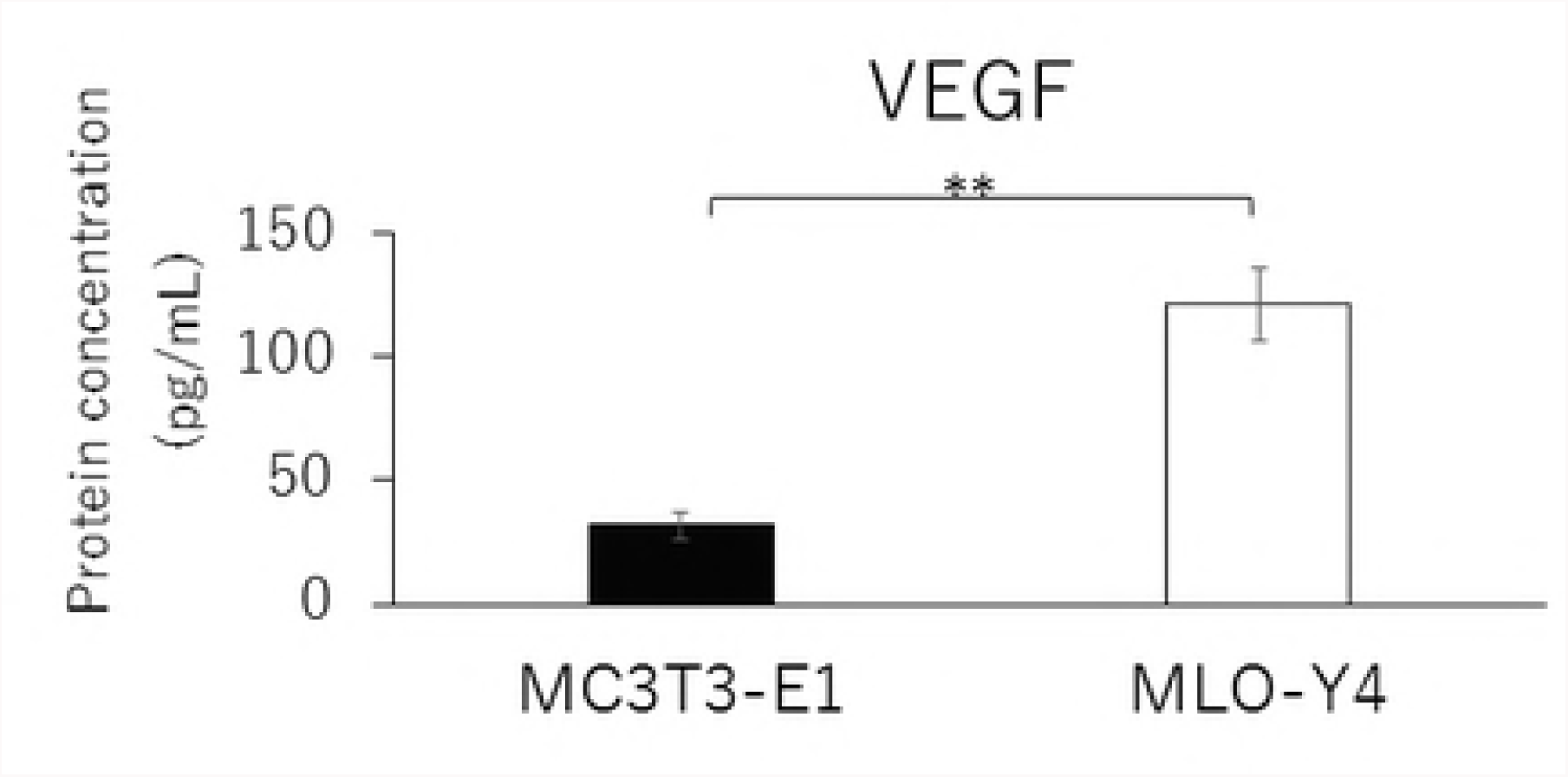
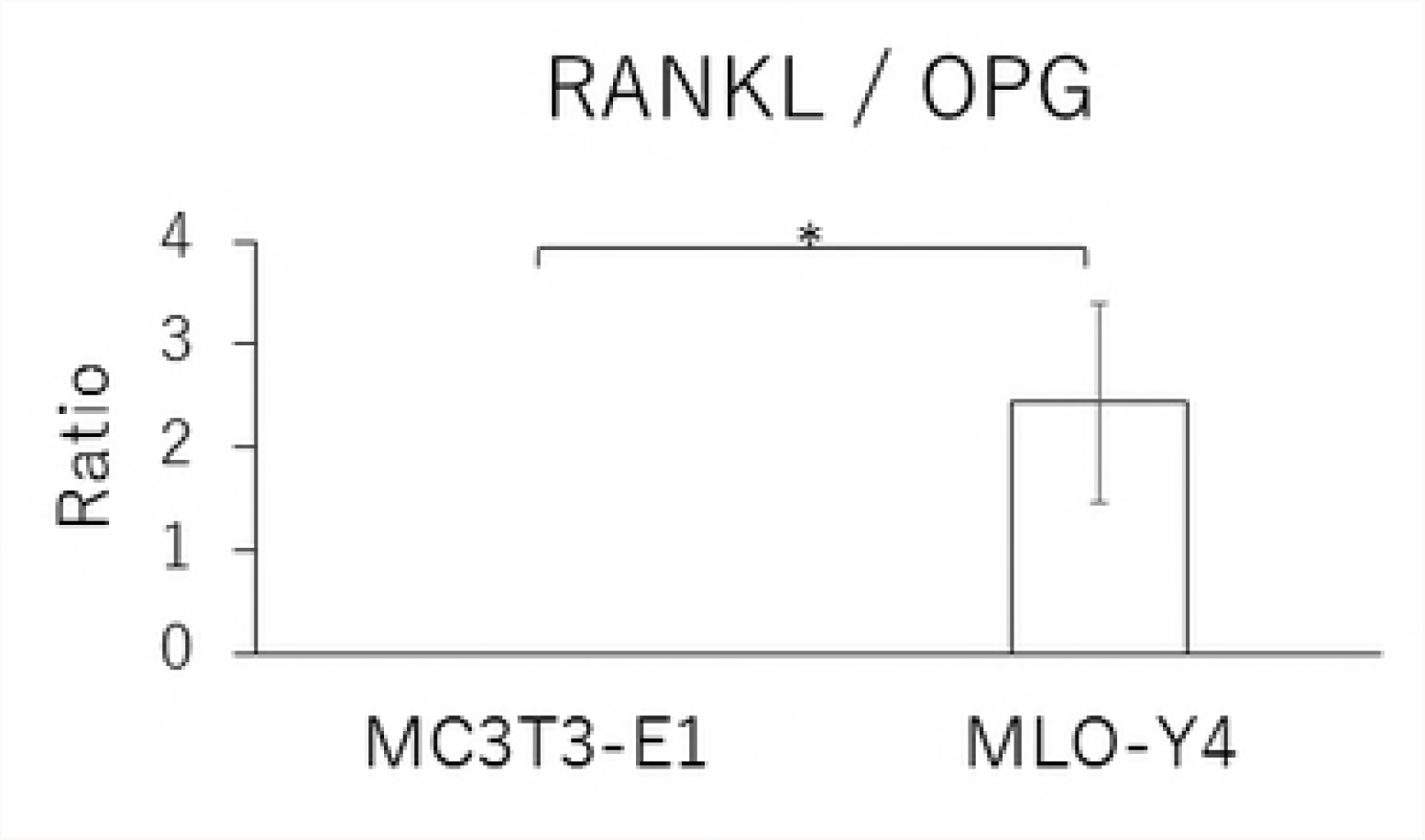
Comparison in the gene expression of RANKL (A), OPG (B), VEGF (C) and RANKL / OPG ratio (D), and the protein concentration of RANKL (E), OPG (F), VEGF (G) and RANKL / OPG ratio (H) between MC3T3-E1 and MLO-Y4 cells. (**; P<0.01)

Similarly, MLO-Y4 cells had a significantly lower level of OPG secretion (Fig. 2F), but higher levels of RANKL and VEGF secretions as well as a higher RANKL/OPG ratio (Figs. 2E, 2G, and 2H) compared with MT3T3-E1 cells.

### Time-course effects of 1.0 g/cm² CF on the expressions of RANKL, OPG, and VEGF as well as the RANKL/OPG ratio

MC3T3-E1 and MLO-Y4 cells were cultured with or without 1.0 g/cm^2^ CF for up to 12 hours. RANKL gene expression and the RANKL/OPG ratio reached maximum levels after 3 hours of CF application in both MC3T3-E1 and MLO-Y4 cells (Figs. 3A, 3D, 3E, and 3H). VEGF mRNA levels in MLO-Y4 cells were maximum 3 hours after CF application (Fig 3G). Protein levels of RANKL and VEGF in MC3T3-E1 cells of the CF group were significantly higher than that of the control group at 3, 6, and 12 hours (Fig 4A); and 6 and 12 hours; respectively (Fig 4C). Protein levels of RANKL and VEGF in MLO-Y4 cells of the CF group were significantly higher than that of the control group at 6 and 12 hours (Fig 4E); and 1, 3, and 12 hours; respectively (Fig 4G). Both MC3T3-E1 and MLO-Y4 cells showed a significant increase in the RANKL/OPG ratio at 3, 6, and 12 hours (Figs 4D and 4H).

**Figs 3.**
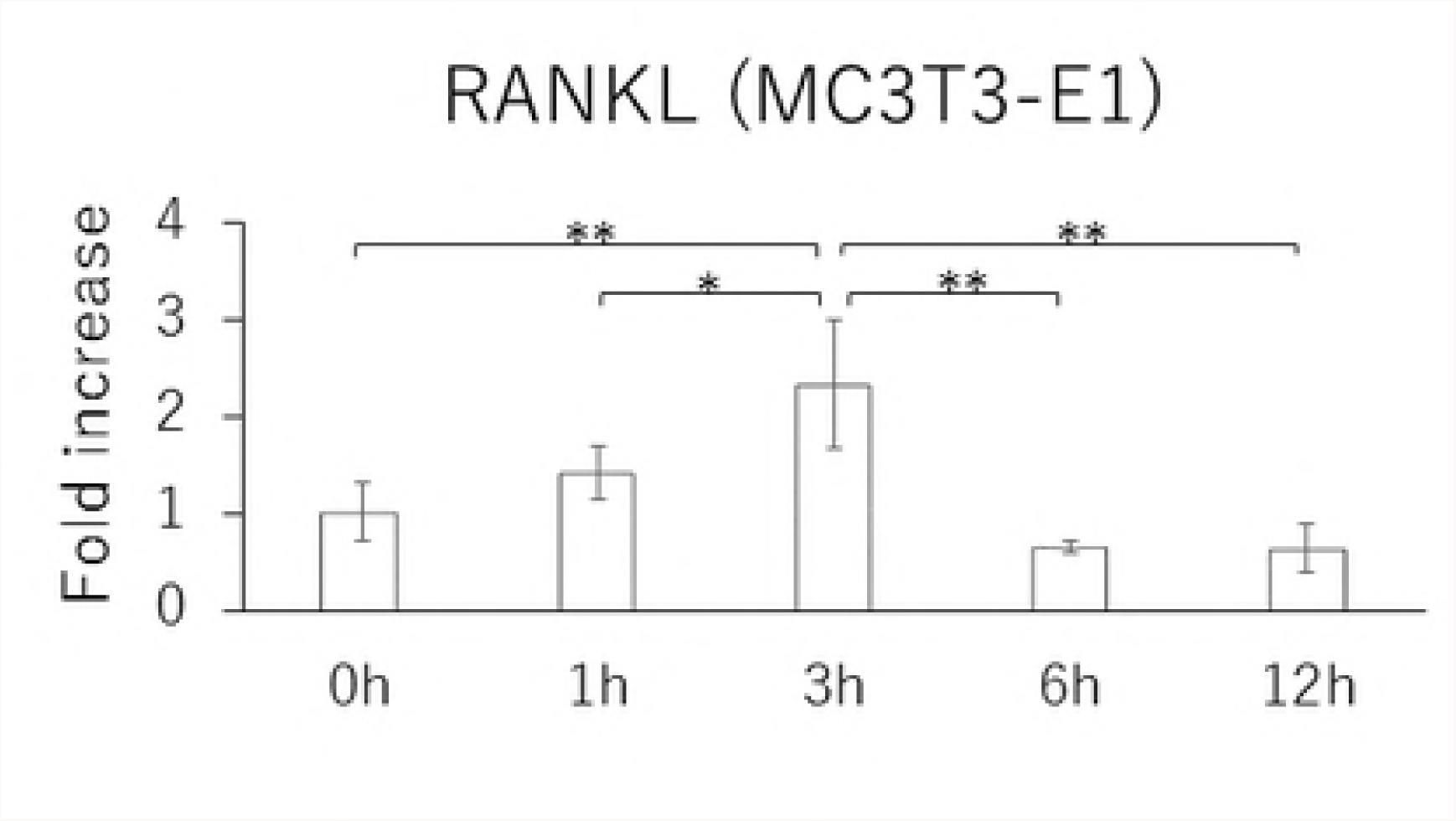
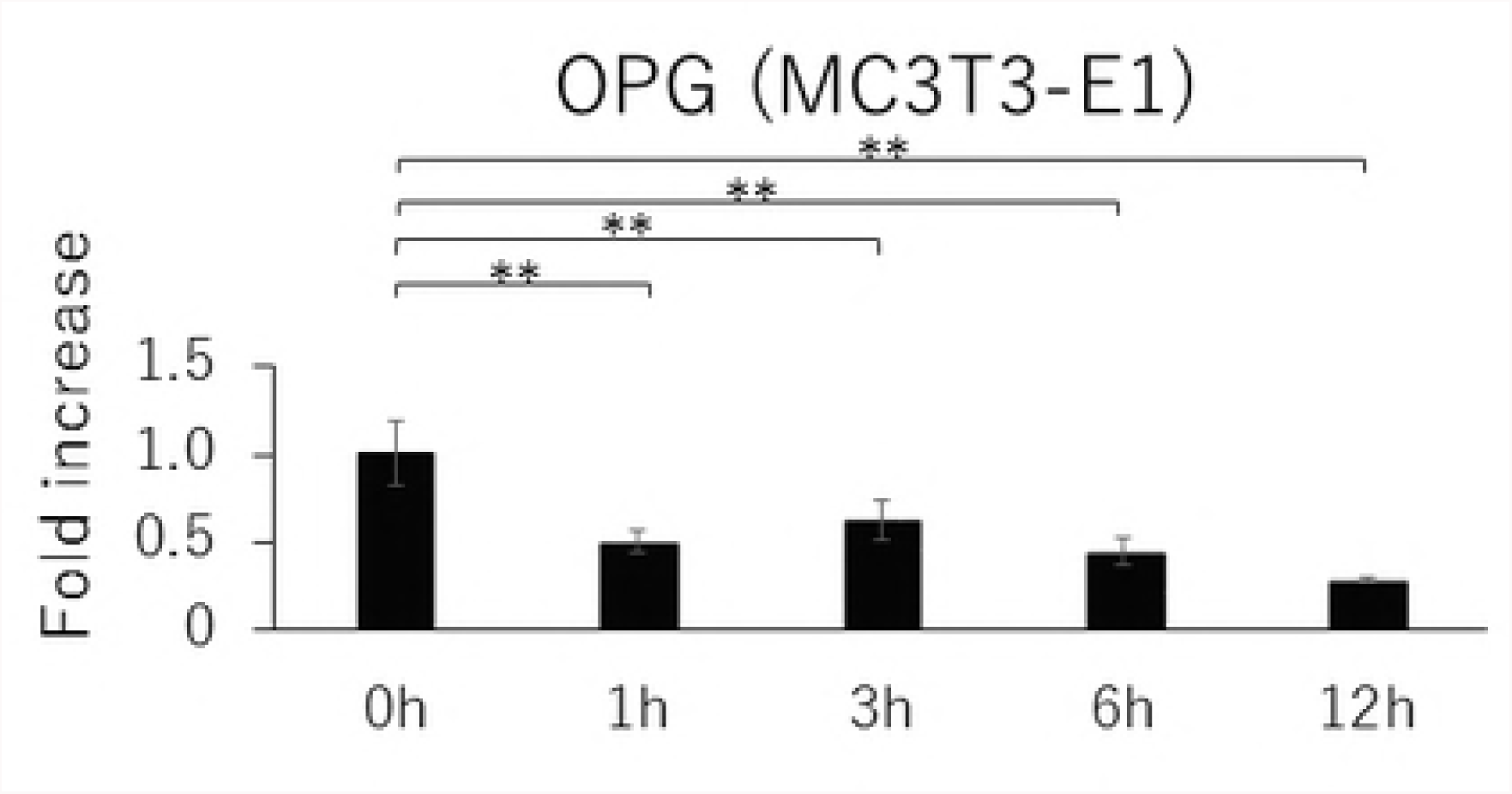
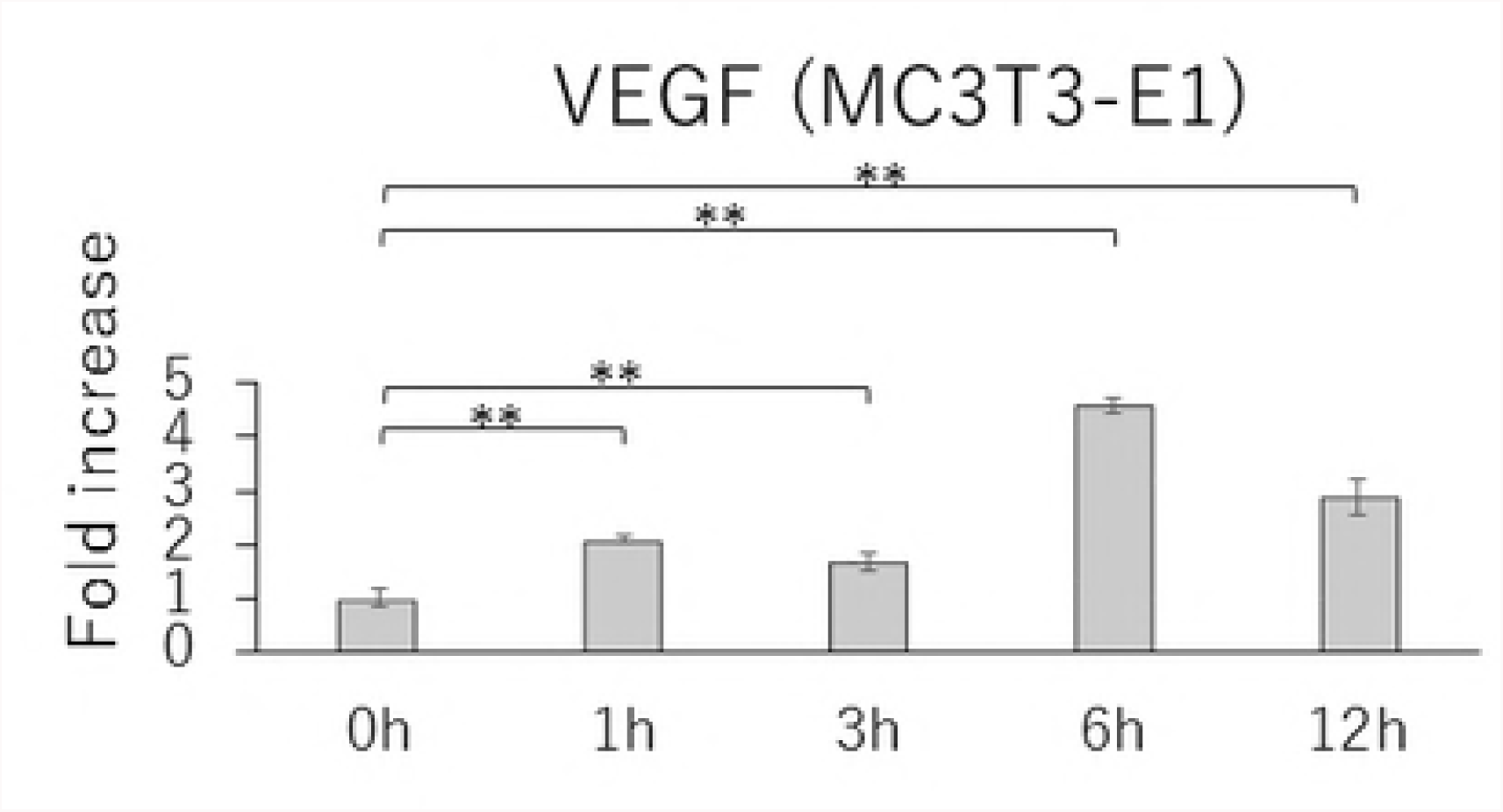
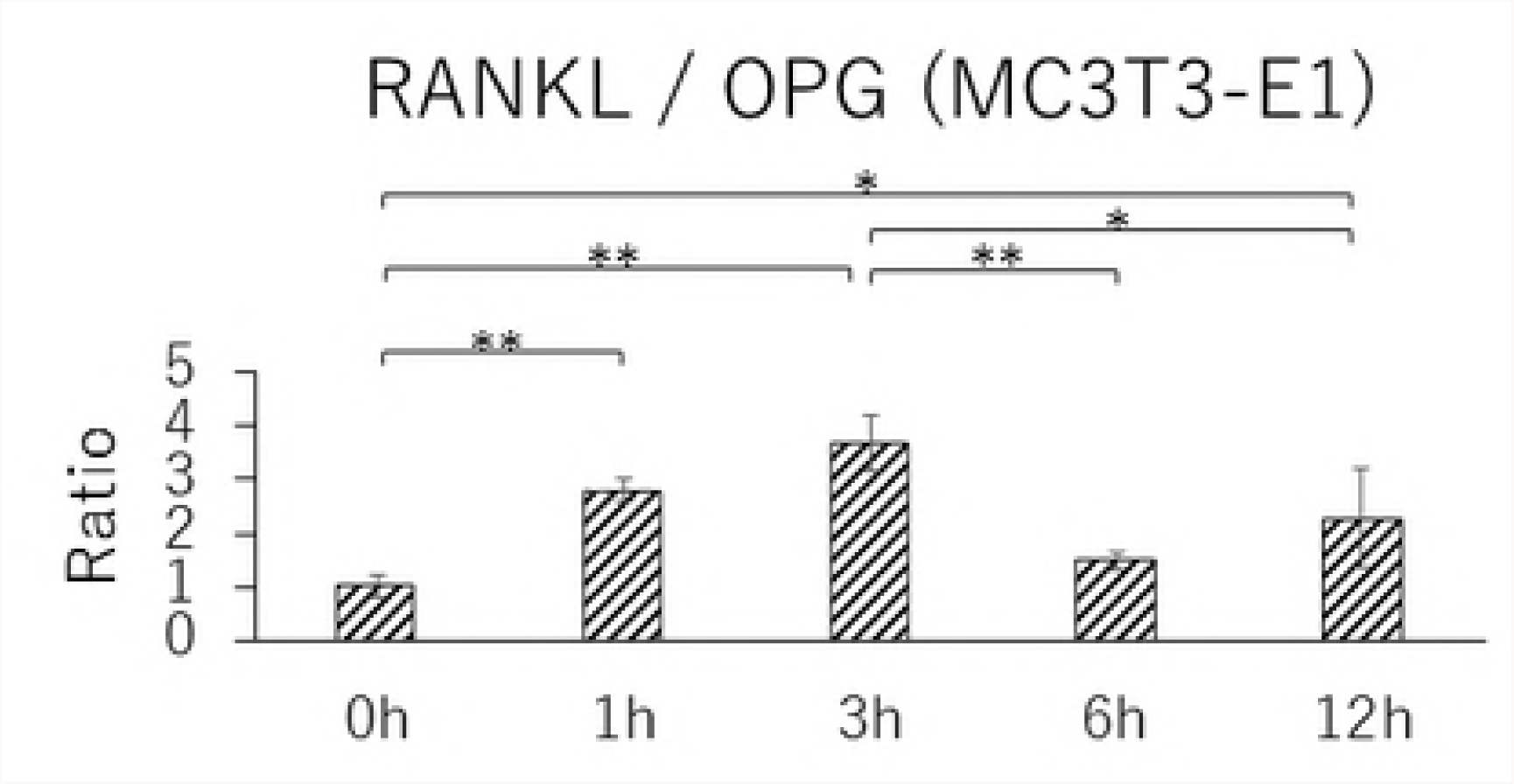
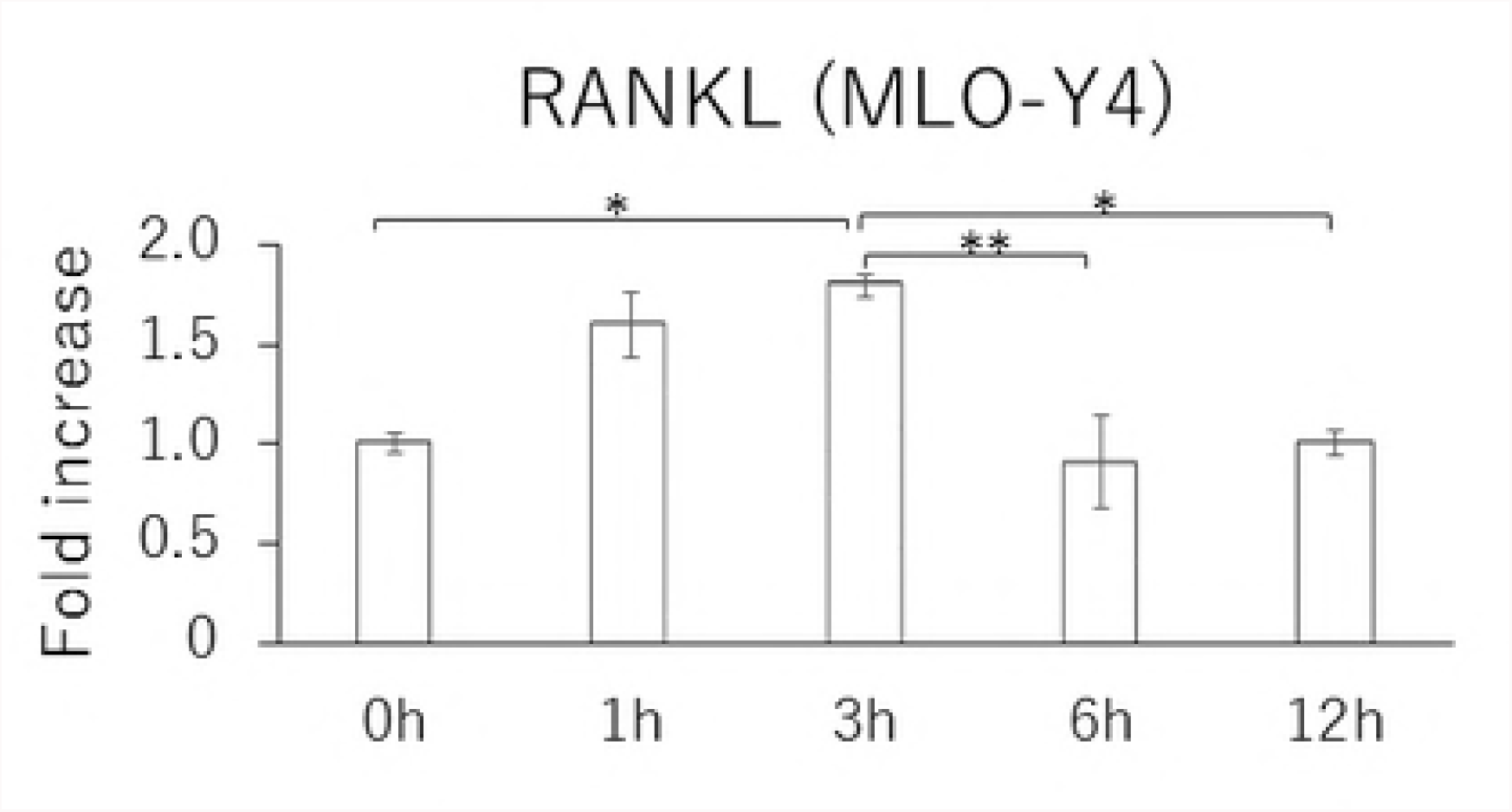
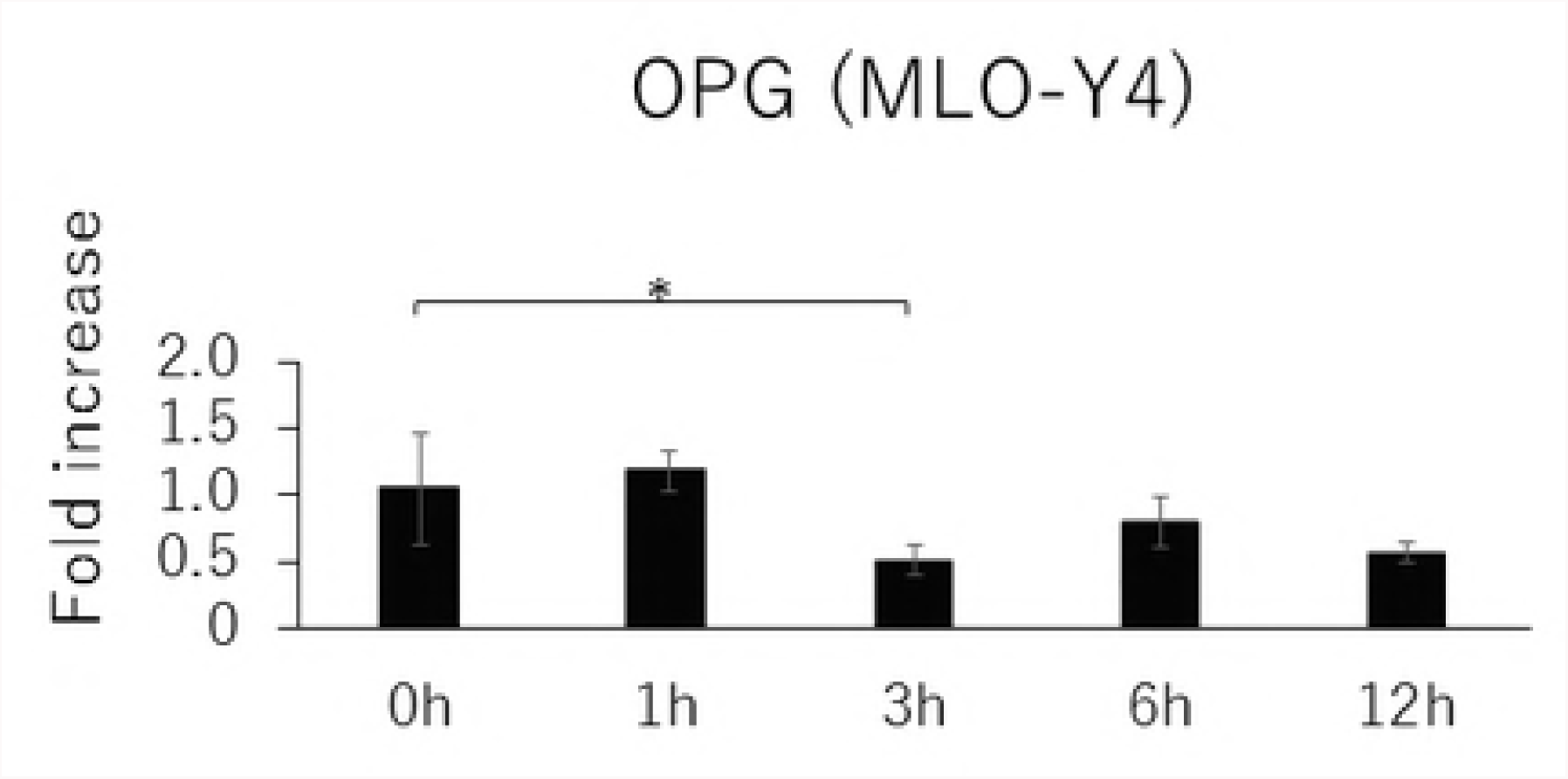
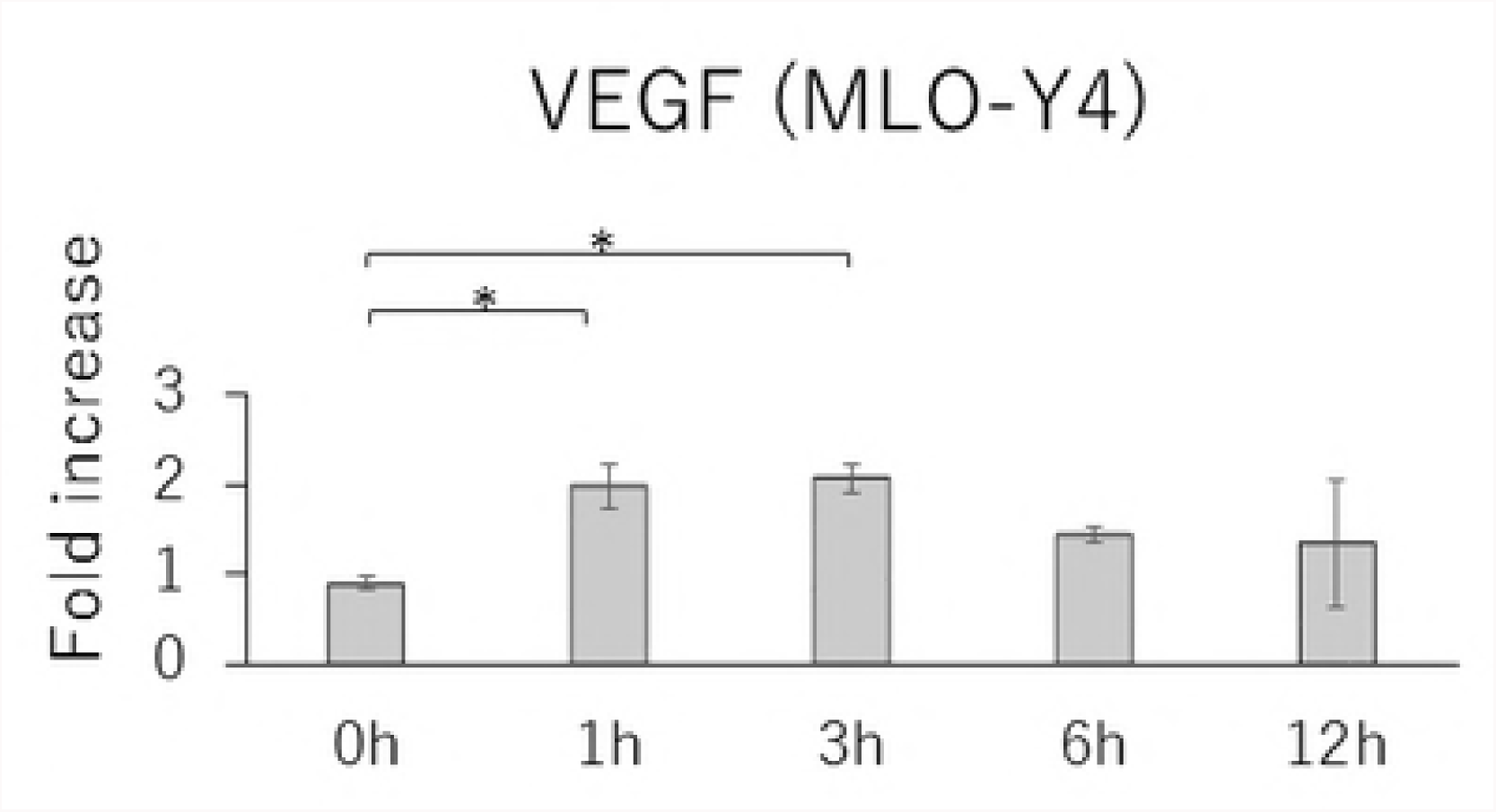
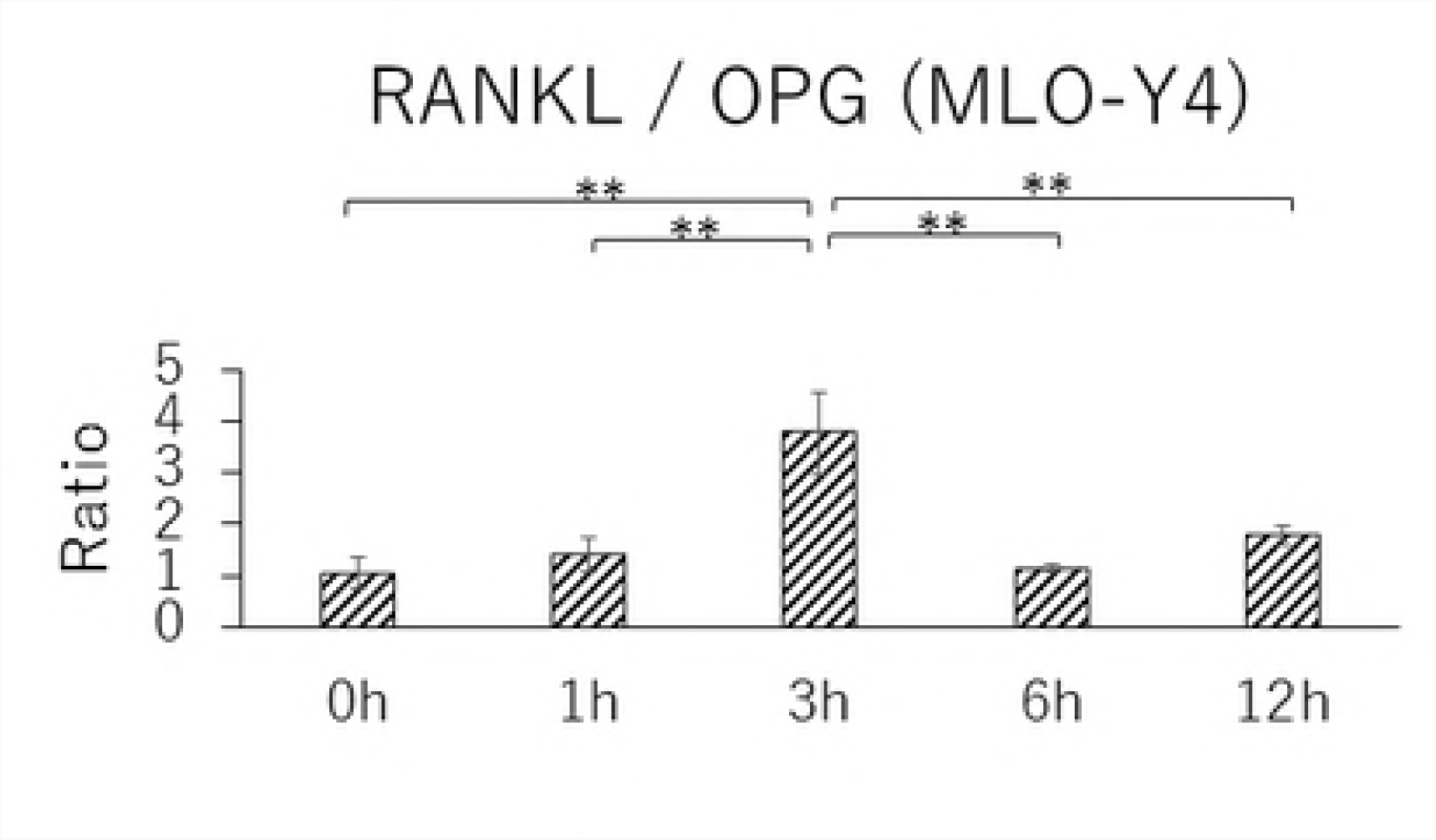
Changes in the gene expression of RANKL (A), OPG (B), VEGF (C) and RANKL / OPG ratio (D) in MC3T3-E1 cells and RANKL (E), OPG (F), VEGF (G) and RANKL / OPG ratio (H) in MLO-Y4 cells after 1, 3, 6 and 12 hours CF application of 1.0 g / cm^2^. (*; P<0.05 and **; P<0.01 respectively)

**Figs 4.**
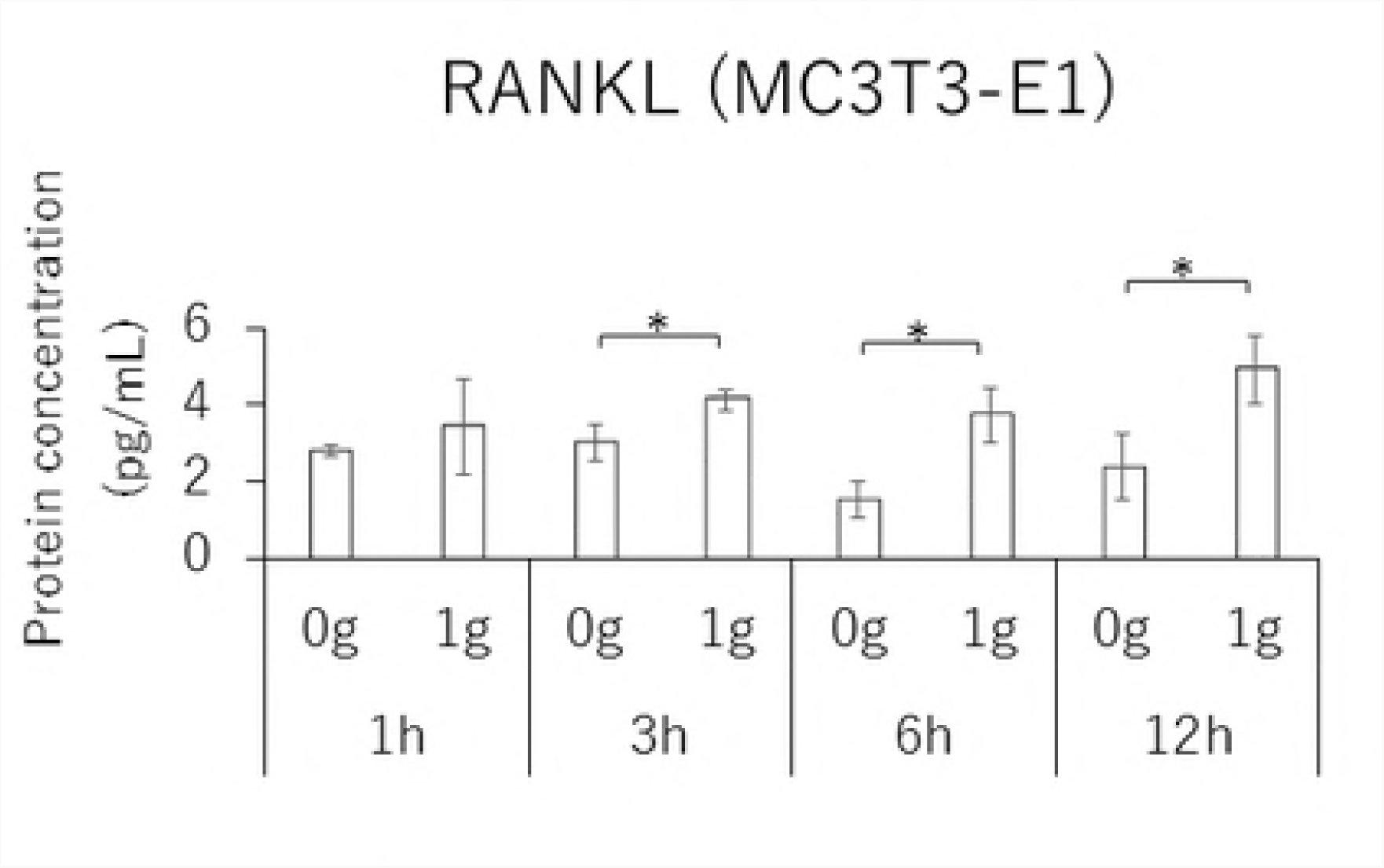
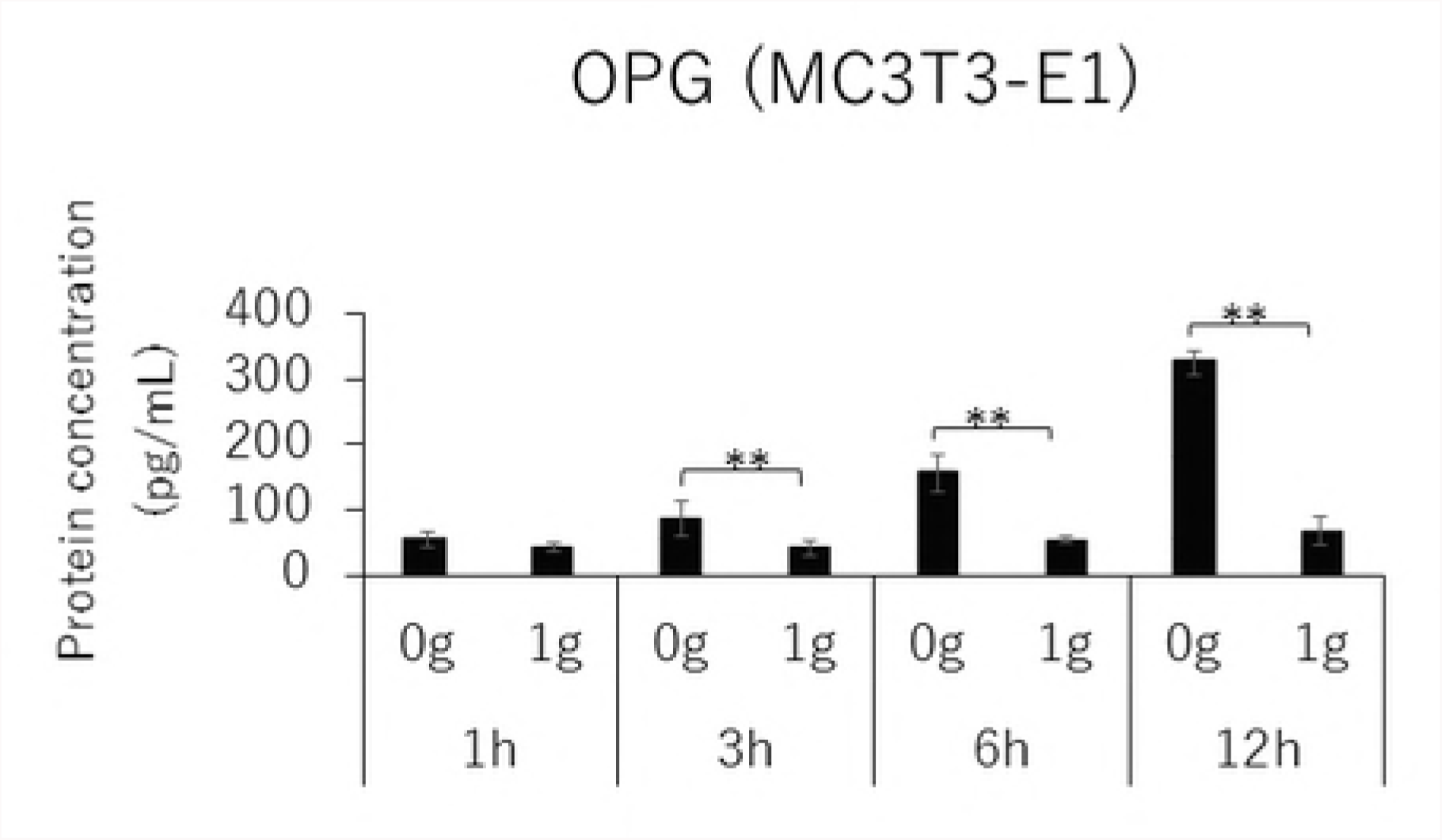
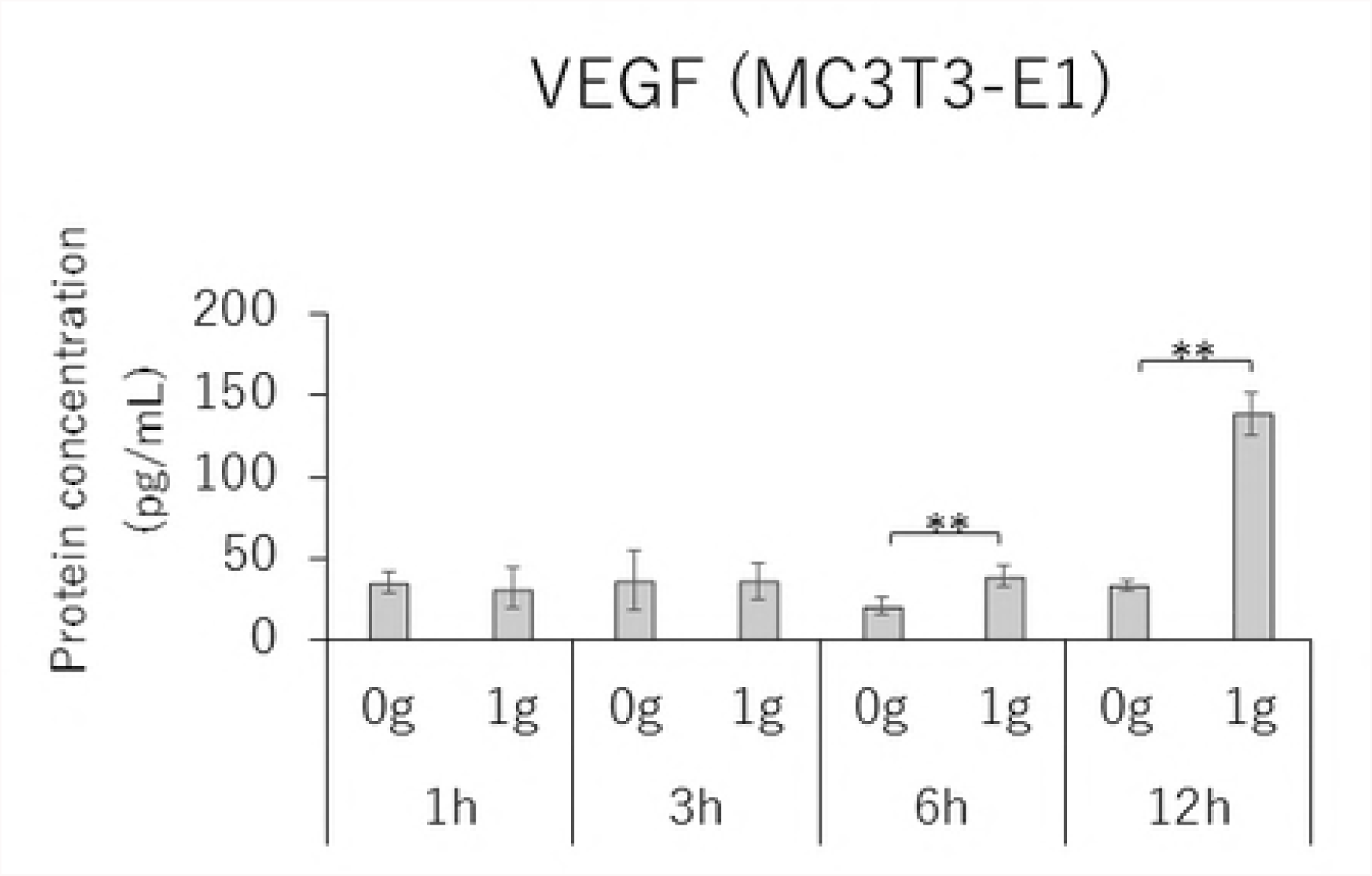
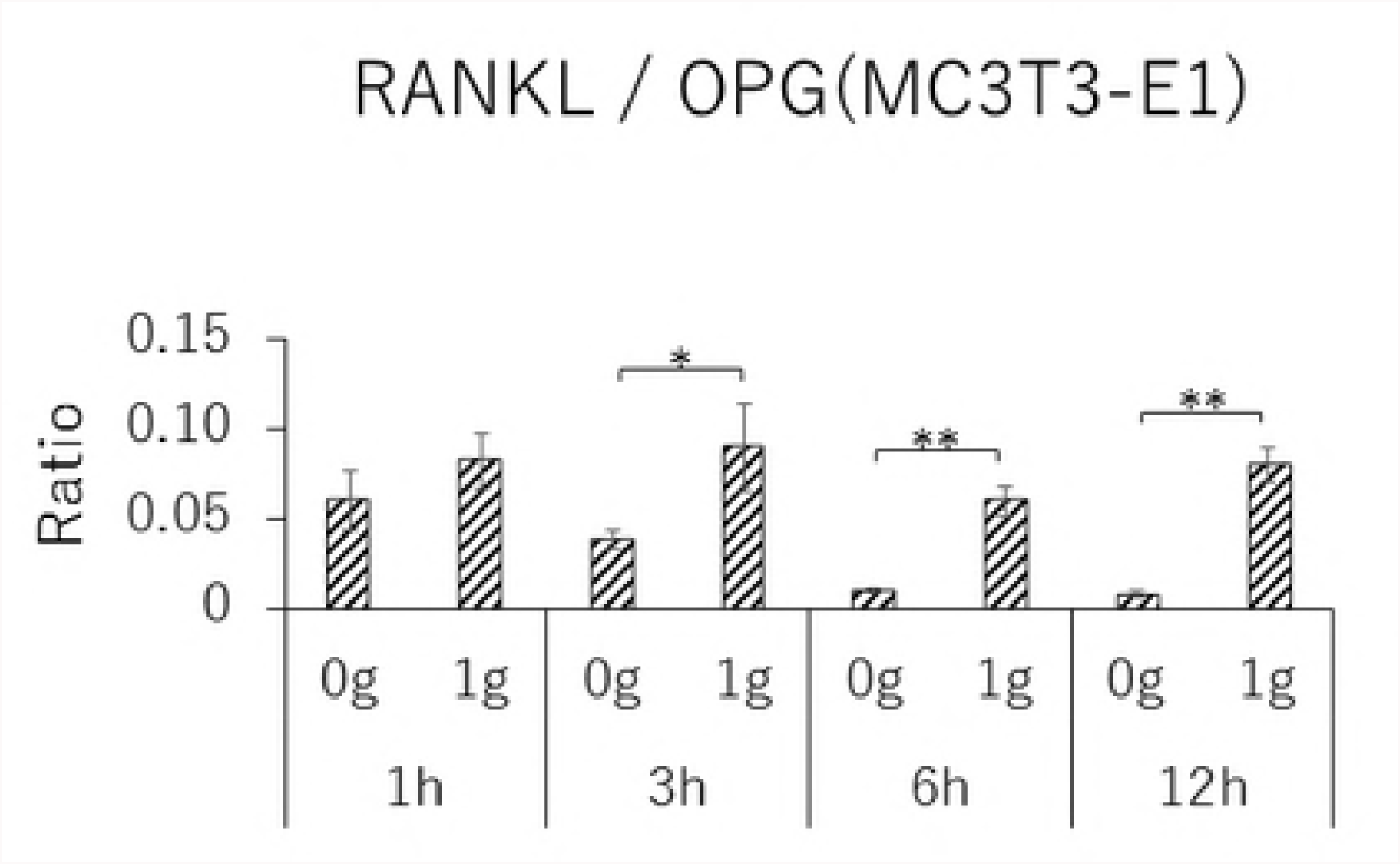
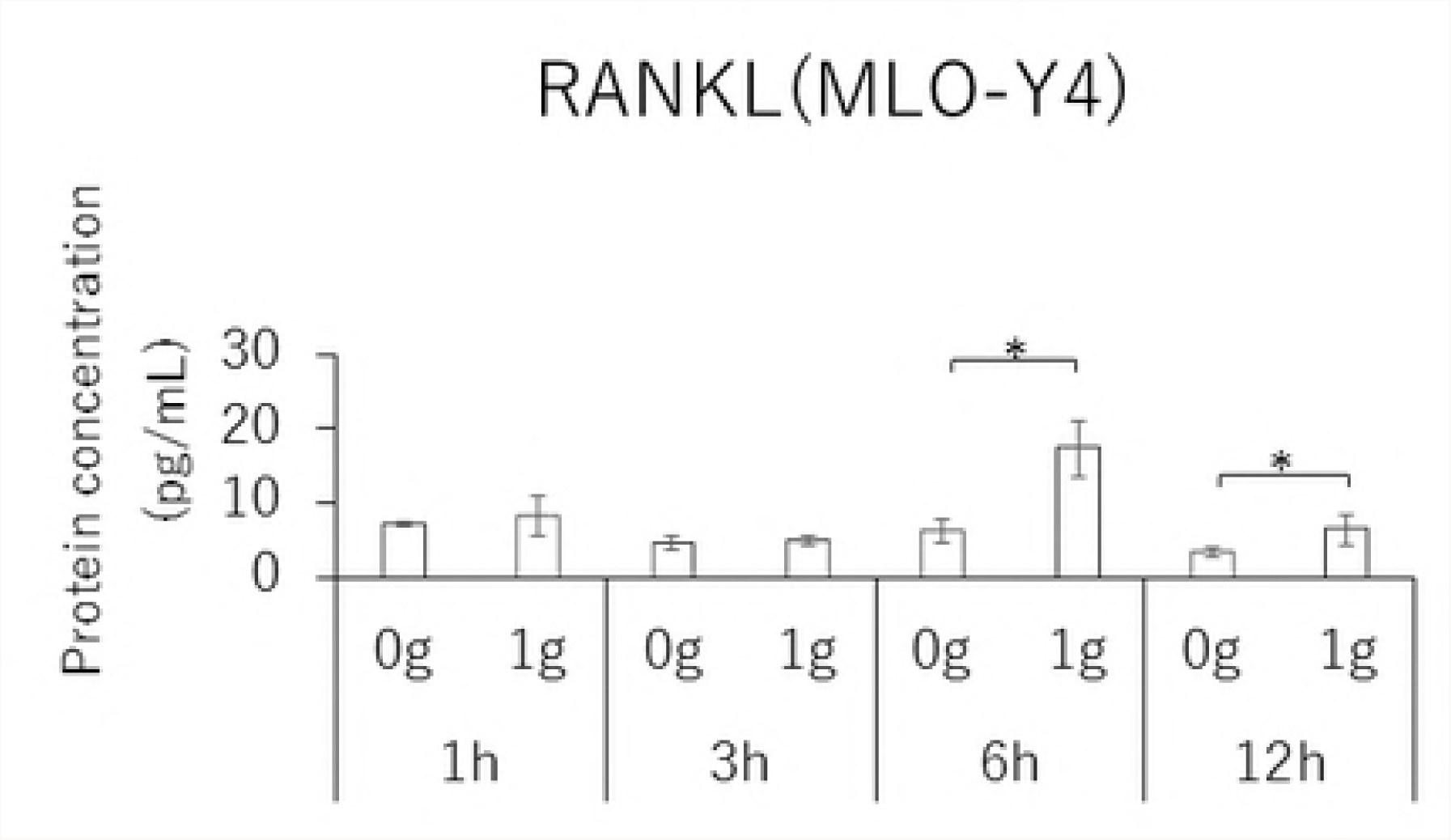
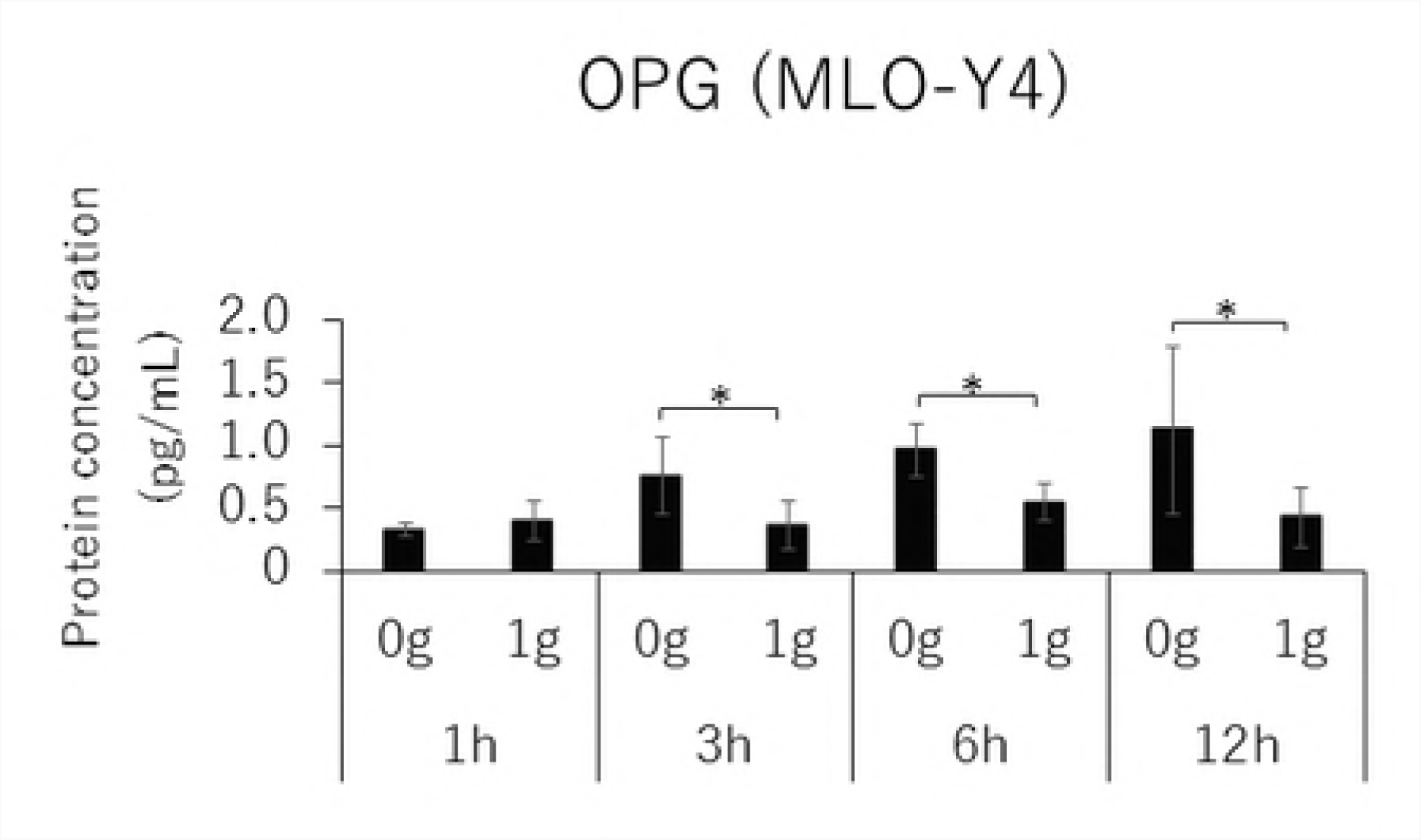
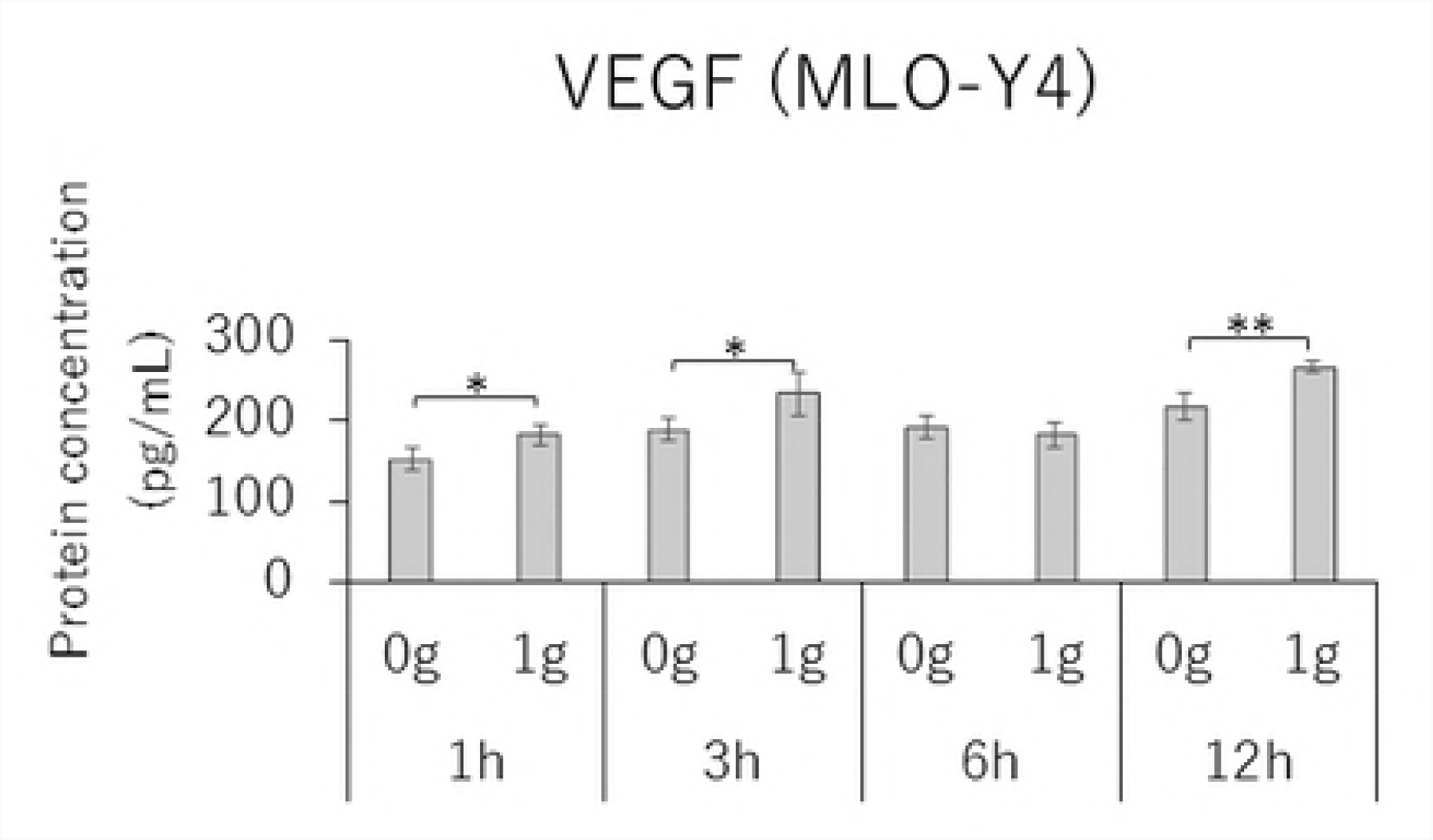
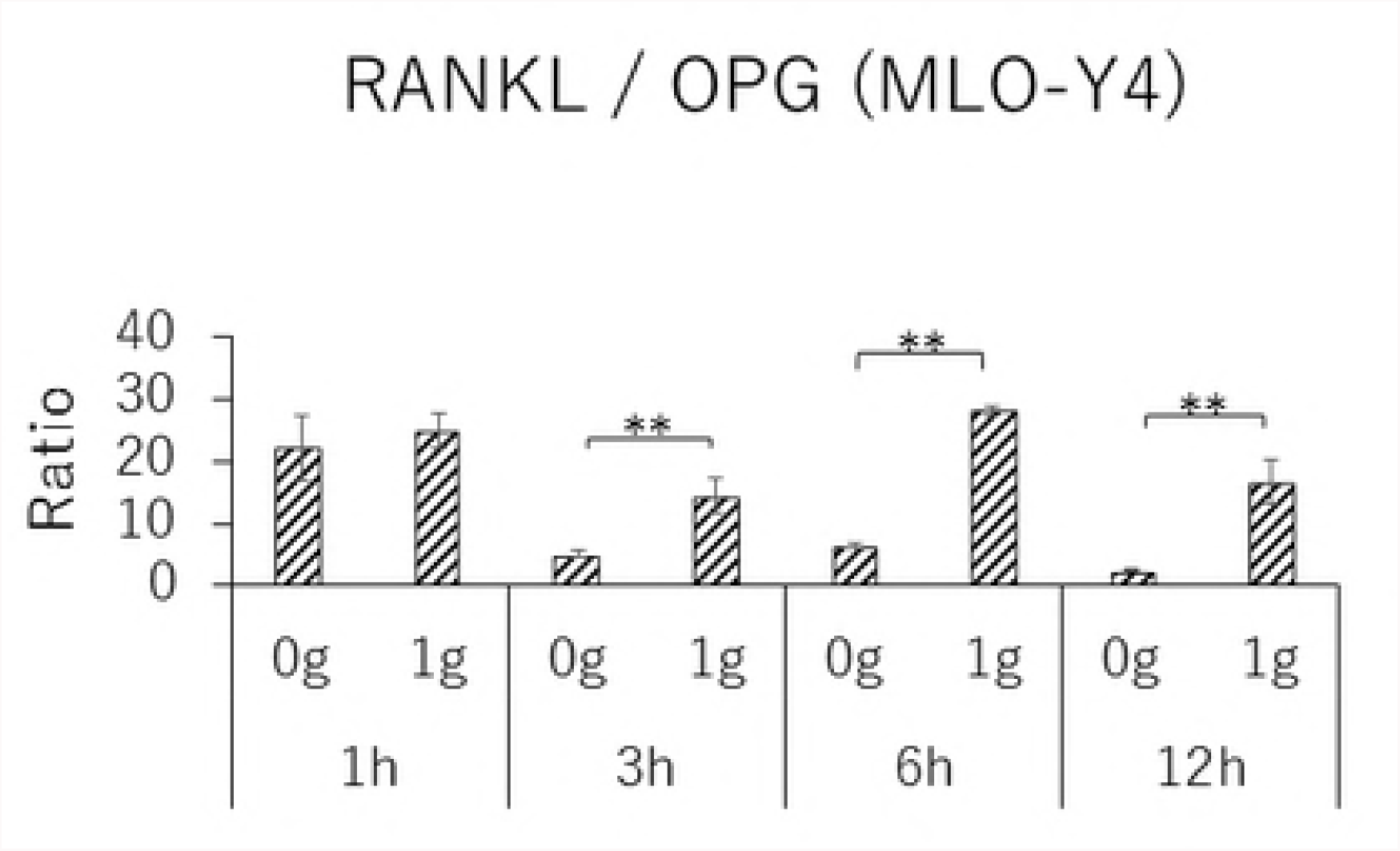
Protein concentration of RANKL (A), OPG (B), VEGF (C) and RANKL / OPG ratio (D) in MC3T3-E1 cells and RANKL (E), OPG (F), VEGF (G) and RANKL / OPG ratio (H) in MLO-Y4 cells after 1, 3, 6 and 12 hours CF application of 1.0 g / cm^2^. (*; P<0.05 and **; P<0.01 respectively)

### Effect of Gd^3+^ Treatment in MLO-Y4 cells

In MLO-Y4 cells, RANKL gene expression and the RANKL/OPG ratio in the CF group were significantly reduced by treatment with 10 μM Gd^3+^ (Figs 5A and 5D). Protein levels of RANKL and VEGF as well as the RANKL/OPG ratio were significantly lower in the Gd^3+^ treatment group than in the non-treatment group (Figs 6A, 6C, and 6D).

**Figs 5.**
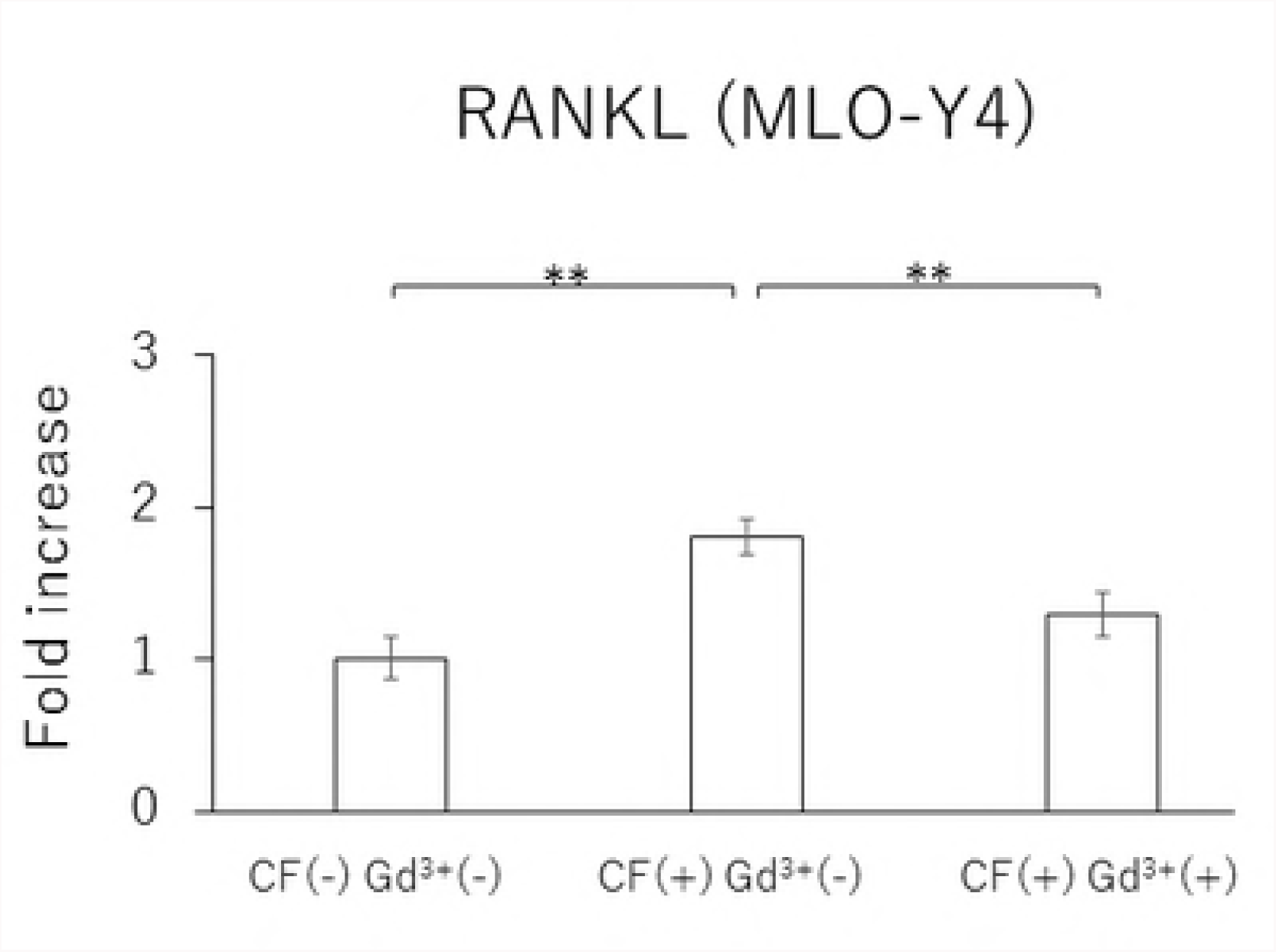
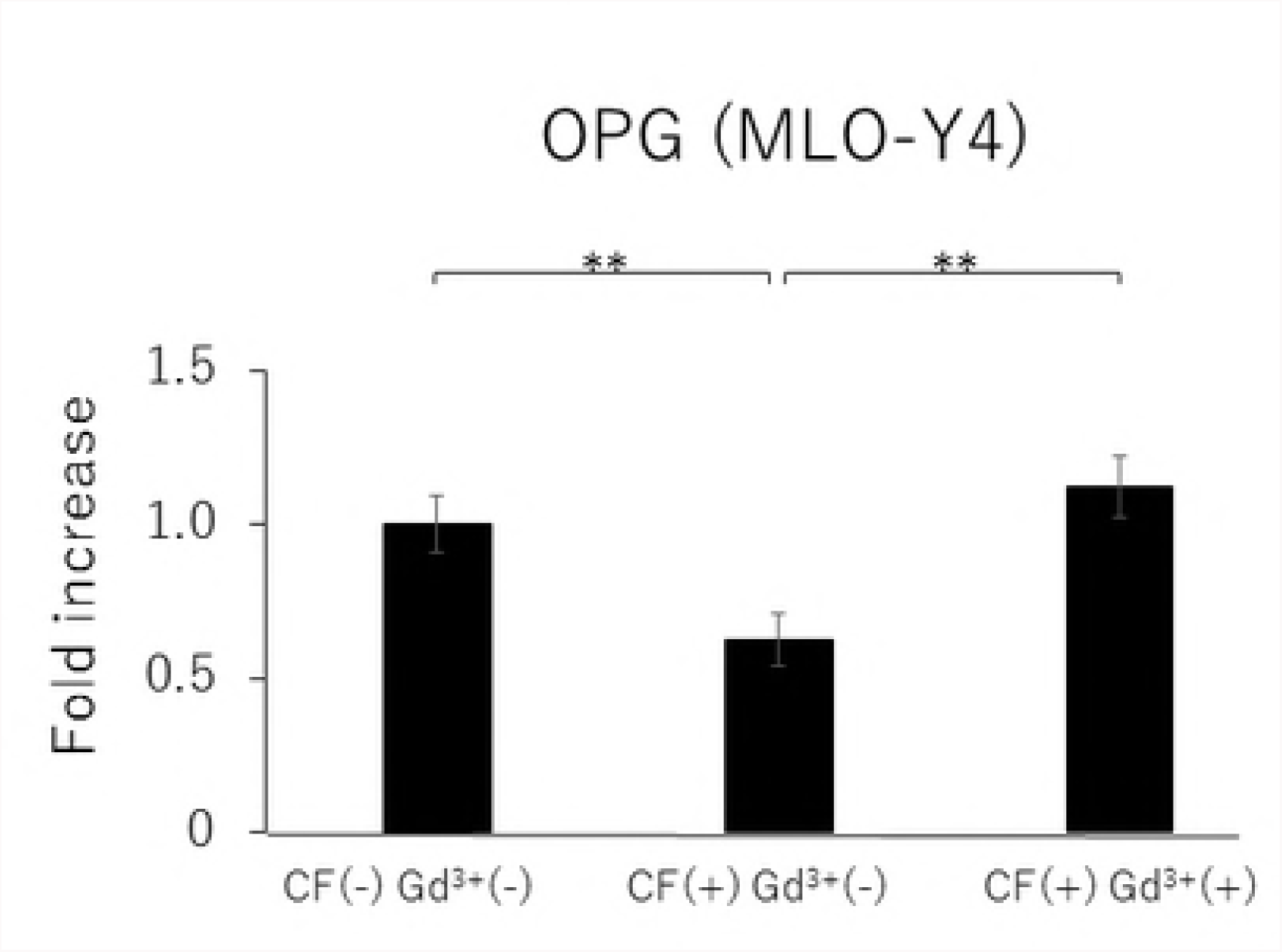
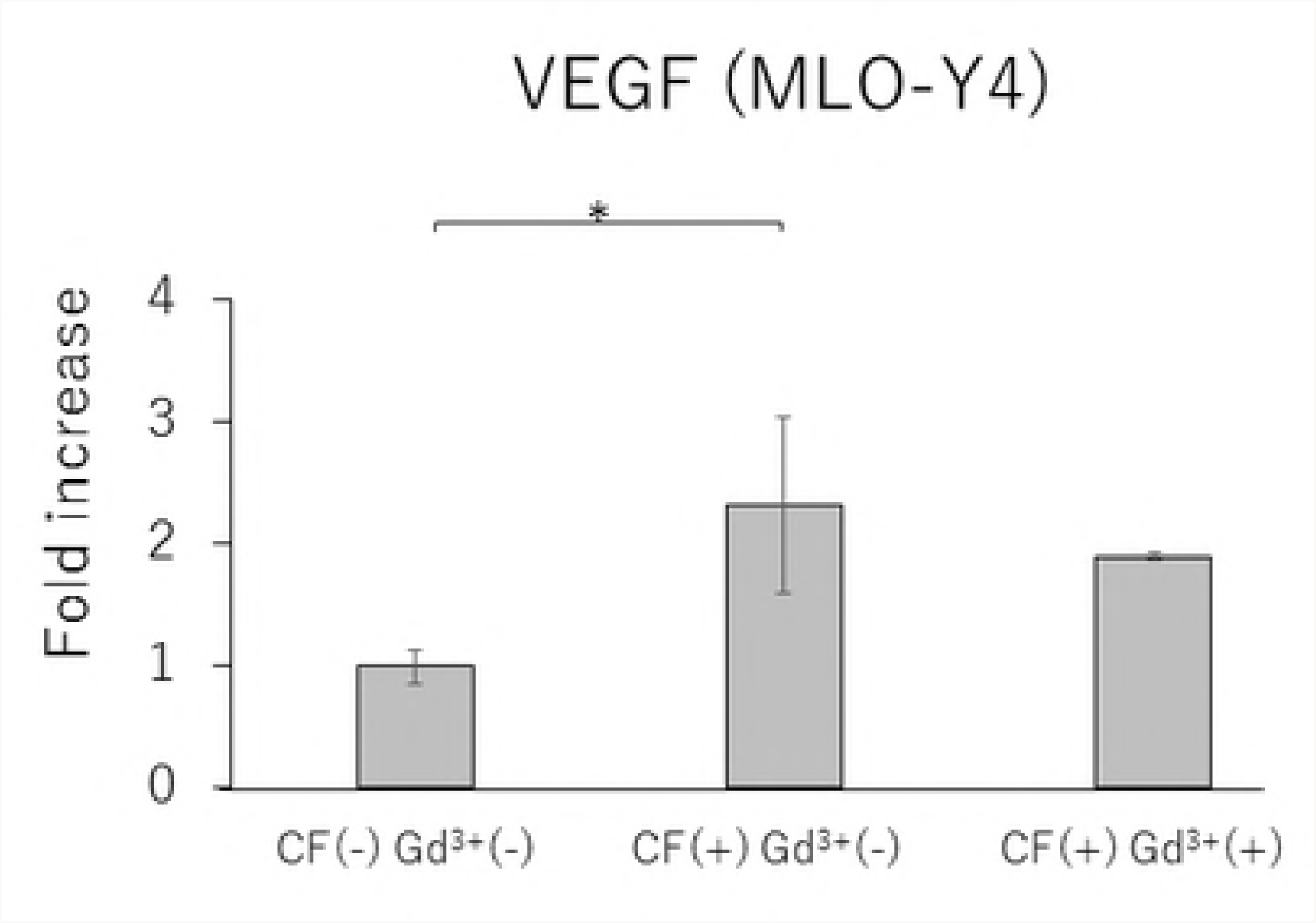
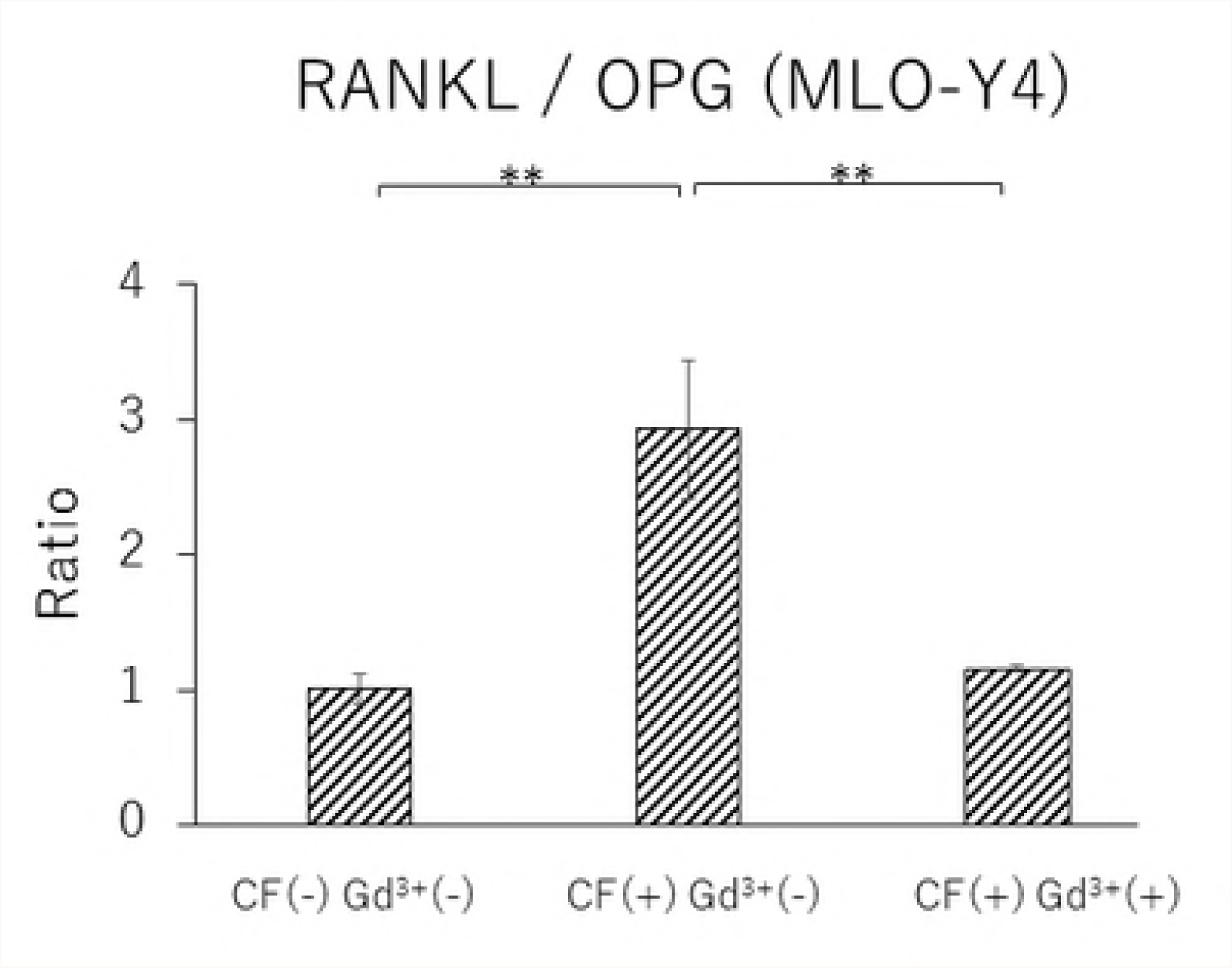
Effects of Gd^3+^ on the gene expression of RANKL (A), OPG (B), VEGF (C) and RANKL / OPG ratio (D) after 1.0 g / cm^2^ CF application in MLO-Y4 cells. (*; P<0.05 and **; P<0.01 respectively)

**Figs 6.**
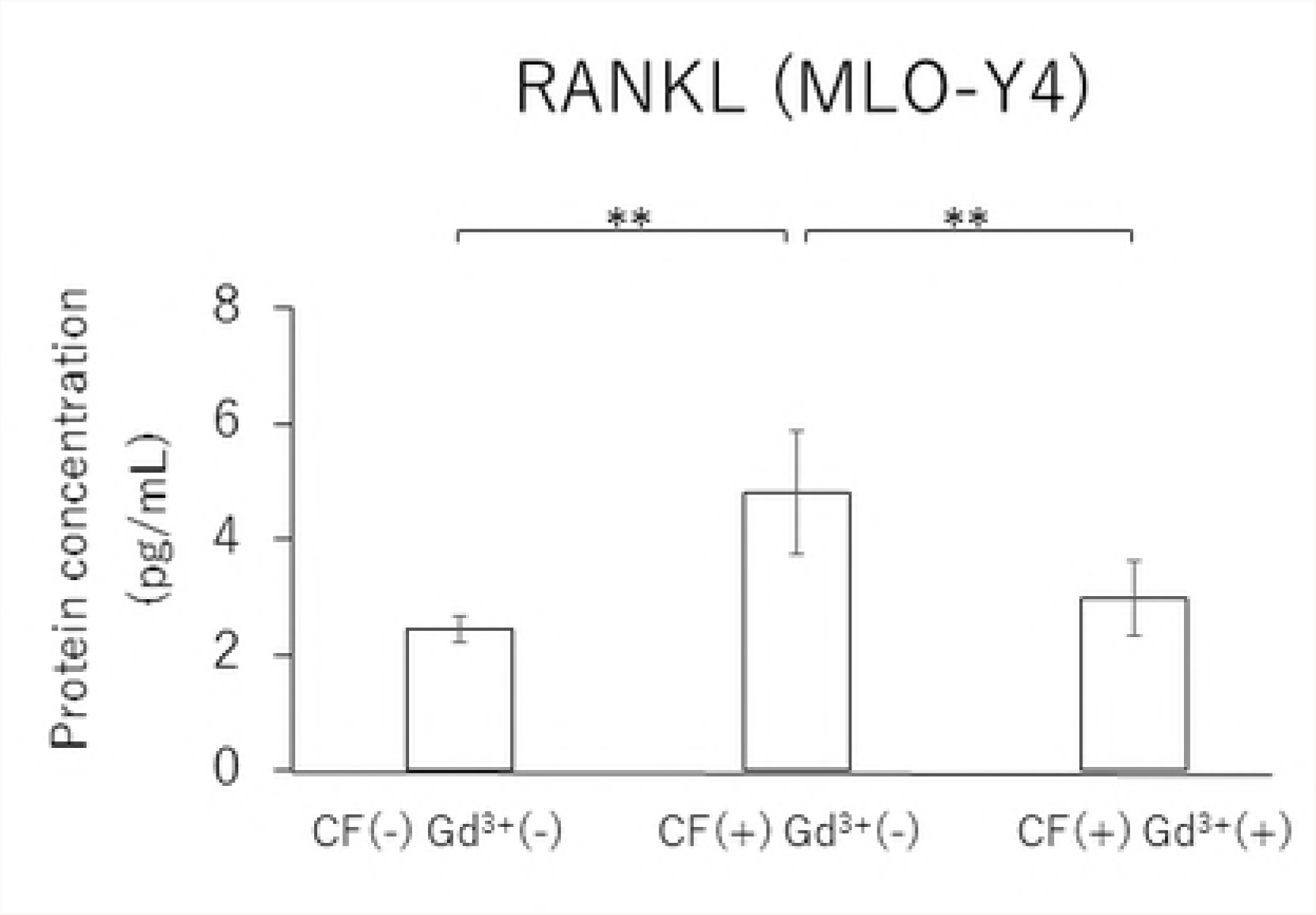
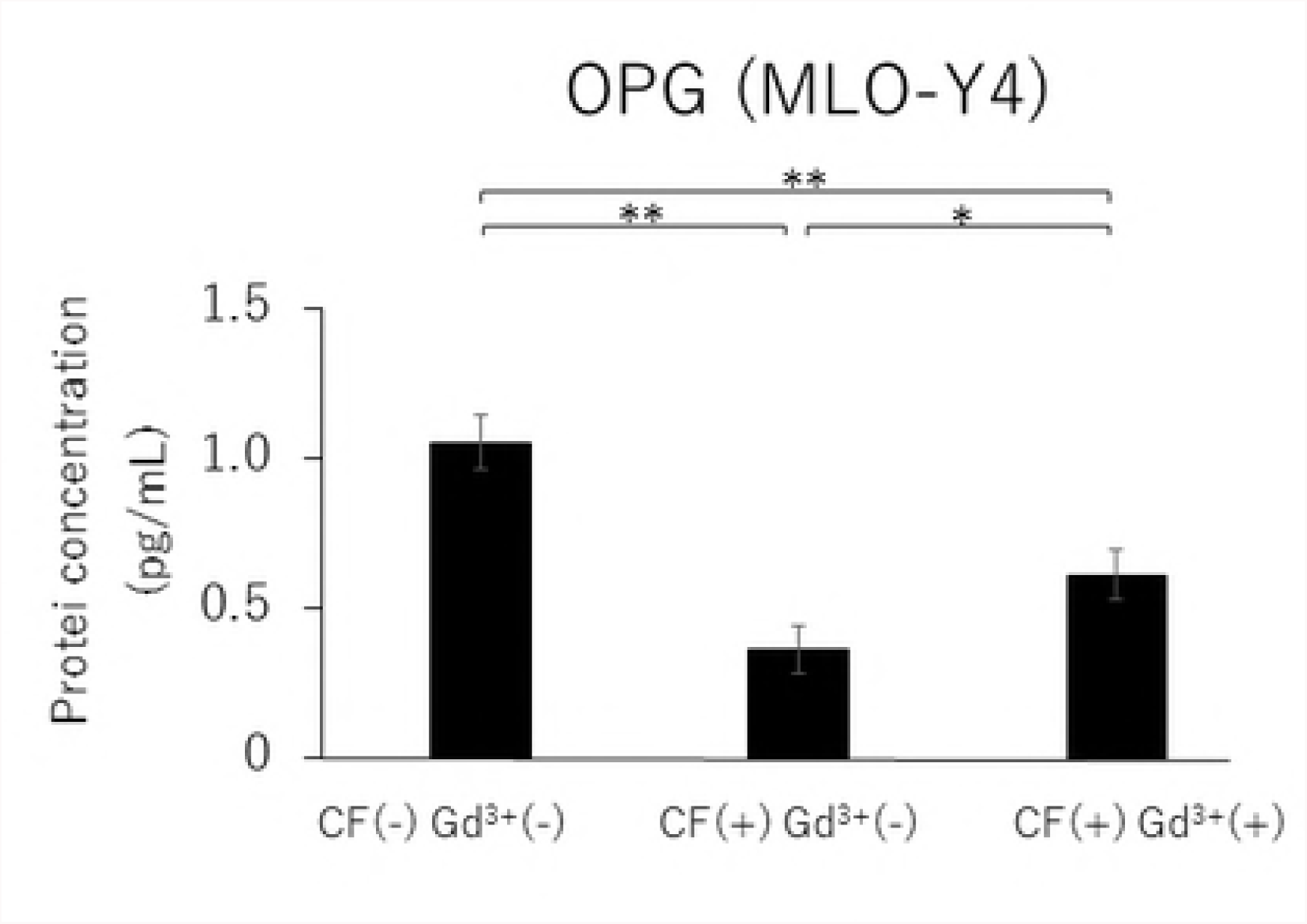
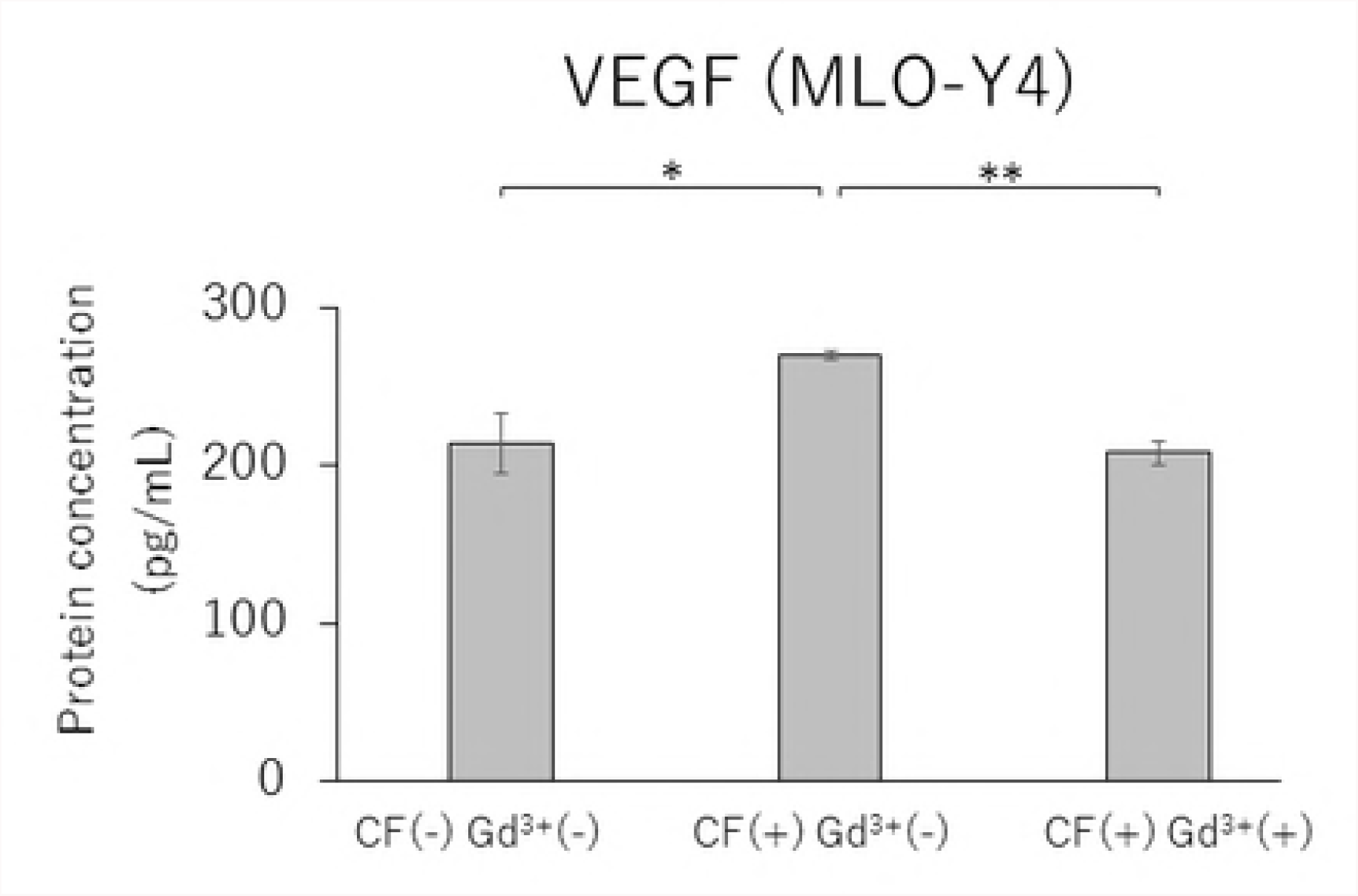
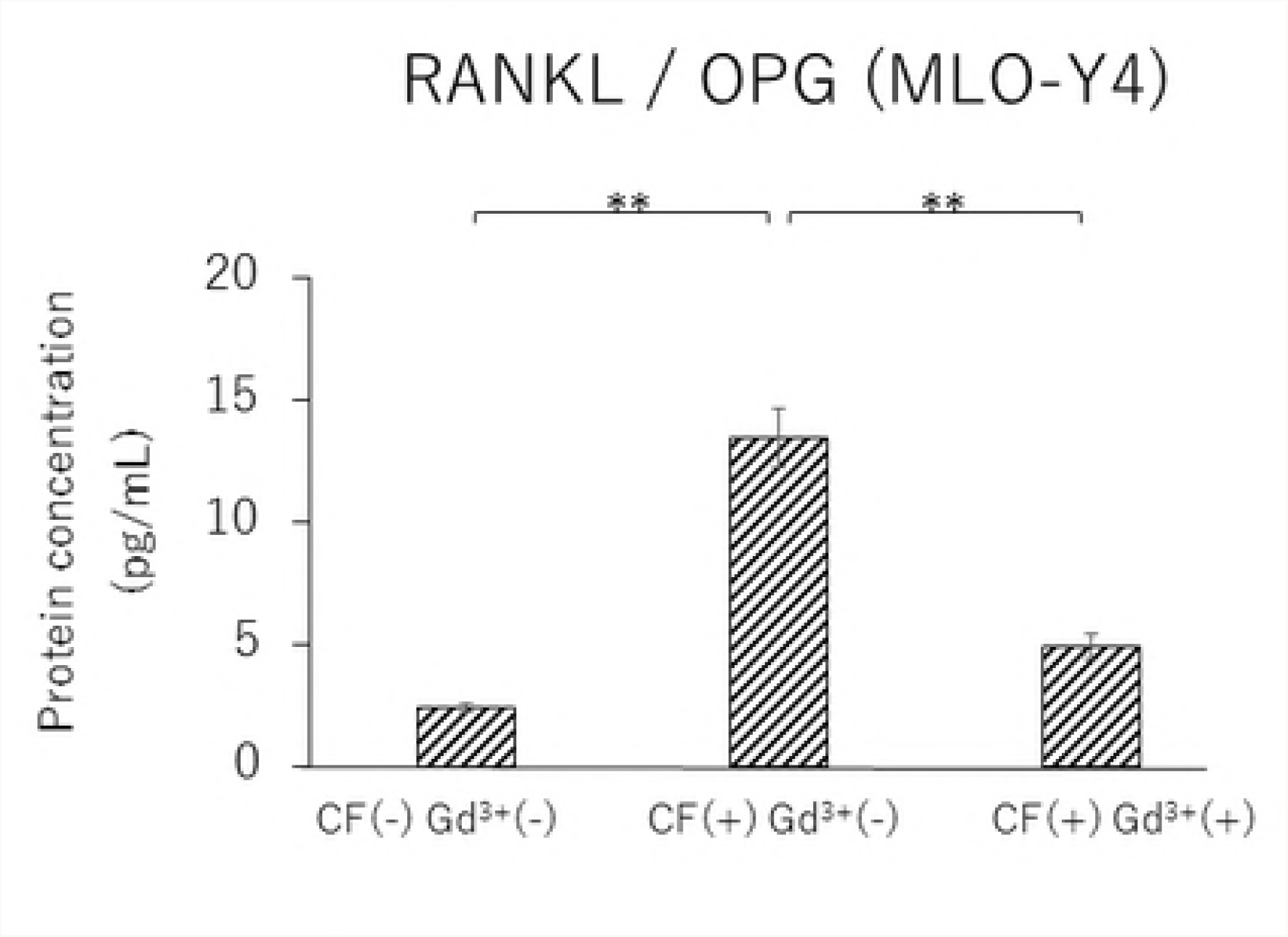
Effects of Gd^3+^ on the protein concentration of RANKL (A), OPG (B), VEGF (C) and RANKL / OPG ratio (D) after 1.0 g / cm^2^ CF application in MLO-Y4 cells. (*; P<0.05 and **; P<0.01 respectively)

## Discussion

Osteoblasts have been considered a main source of RANKL and OPG, which play essential roles in osteoclastogenesis [6, 7]. However, recent studies have demonstrated that osteocytes mainly regulate physiological osteoclastogenesis by the production of RANKL during bone remodeling [18, 19]. In this study, the mRNA and protein expression levels of RANKL as well as the RANKL/OPG ratio in MLO-Y4 cells were significantly higher than in MC3T3-E1 cells, which is in accordance with results from previous studies [18, 19]. Furthermore, the gene and protein expression levels of OPG were significantly lower in MLO-Y4 cells than in MC3T3-E1 cells. These results suggested that osteocytes play a more important role for osteoclastogenesis by enhancing RANKL expression and suppressing OPG expression compared to osteoblasts.

Osteocytes are differentiated from osteoblasts, which are embedded in the bone matrix with extended dendritic processes. For crosstalk among the bone component cells, dendritic processes are thought to be essential because osteocytes connect each other and adjacent cells [17]. Although the importance of cellular reaction for bone remodeling is well known, the mechanism that osteocytes use to promote cytokines for bone remodeling after receiving mechanical stress, especially CFs, is still unclear. Nettelhoff et al. reported that the mRNA expression of RANKL as well as the RANKL/OPG ratio peaked in osteoblasts after 5% CF application by using the Flexcell Compression Plus System [12]. Sanchez et al. also used the Flexcell Compression Plus System, and showed that OPG gene expression was significantly decreased after 4 hours of CF in osteoblasts [22]. Tripuwabhrut et al. reported that the application of 4.0 g/cm² CF induced significantly increased expression of RANKL mRNA in osteoblasts. They also showed that 2.0 and 4.0 g/cm² CF significantly reduced the expression of OPG mRNA [13]. In this study, gene and protein expression levels of RANKL in MLO-Y4 and MC3T3-E1 cells reached a maximum 3 hours after applying a CF of 1.0 g/cm^2^. The application of a CF of 1.0 g/cm² induced a significant decrease of OPG gene and protein expressions both in MLO-Y4 and MC3T3-E1 cells. From these findings, both osteocytes and osteoblasts can respond to CF by upregulating RANKL and downregulating OPG.

Angiogenesis is essential for the development and regeneration of various tissues. Especially in bone remodeling, bone resorption by osteoclasts and neovascularization by blood vessel invasion are required for bone morphogenesis and growth. VEGF is a mitogen specific to vascular endothelial cells and promotes angiogenesis [10]; VEGF also acts as a vascular permeability factor [23]. Previous studies have shown that recombinant human VEGF can induce osteoclast generation in osteopetrotic (*op/op*) mice, which were characterized by a deficiency in osteoclasts, monocytes, and macrophages due to a lack of functional macrophage-colony stimulating factor (M-CSF) [24, 25]. It was also demonstrated that local administration of rhVEGF during experimental tooth movement can increase the number of osteoclasts and accelerate the rate of tooth movement [26, 27]. Therefore, it is possible that VEGF is closely related to bone remodeling by inducing osteoclasts as well as angiogenesis. Furthermore, Motokawa et al. reported that cyclic tensile forces enhance the mRNA and protein expressions of VEGF in MC3T3-E1 cells. Furthermore, Gd^3+^ treatment has been reported to decrease the amount of VEGF mRNA and protein concentration through the S-A channel [14]. The S-A channel is a membrane stretch-activated ionic channel that was discovered in tissue-cultured embryonic chick skeletal muscle [28]. It was reported that stretched cellular membranes increased intracellular Ca^2+^ concentration in human umbilical endothelial cells. The Ca^2+^ increase was inhibited by administration of Gd^3+^, which is a potent blocker for the S-A channel [15]. These findings suggested that cells can receive mechanical stress mediated by Ca^2+^-permeable S-A channels that exist on the cell membrane. Additionally, 10 μM Gd^3+^ reduced the RANKL and VEGF mRNA and protein expression levels as well as the RANKL/OPG ratio, which was enhanced by 1.0 g/cm^2^ CF application in MLO-Y4 cells. Qin et al. reported that there was a correlation between the degrees of extension of the cell membrane due to the application of CF and the activation of the S-A channel [29], suggesting that osteocytes receive CFs via the S-A channel.

Because the mRNA and protein expression levels of RANKL and VEGF and the RANKL/OPG ratio in MLO-Y4 cells were significantly higher than those in MC3T3-E1 cells in this study, it is possible that ostesocytes might play a more important role in bone metabolism and angiogenesis than osteoblasts, as osteocytes regulate the expression of RANKL, OPG, and VEGF via the S-A channel by responding to mechanical stress.

## Conclusions

The gene and protein expression of RANKL/OPG ratio and VEGF in MLO-Y4 cells showed significantly higher than in MC3T3-E1 cells. After CF application, both cells showed significant increases in RANKL/OPG ratio and VEGF. The upregulated gene and protein levels of these factors were reduced by Gd^3+^ administration.

## Acknowledgements

Editorial support in the form of Medical Writing was provided by Editage.

